# Distinguishing between parallel and serial processing in visual attention from neurobiological data

**DOI:** 10.1101/383596

**Authors:** Kang Li, Mikiko Kadohisa, Makoto Kusunoki, John Duncan, Claus Bundesen, Susanne Ditlevsen

## Abstract

Serial and parallel processing in visual search have been long debated in psychology but the processing mechanism remains an open issue. Serial processing allows only one object at a time to be processed, whereas parallel processing assumes that various objects are processed simultaneously. Here we present novel neural models for the two types of processing mechanisms based on analysis of simultaneously recorded spike trains using electrophysiological data from prefrontal cortex of rhesus monkeys while processing task-relevant visual displays. We combine mathematical models describing neuronal attention and point process models for spike trains. The same model can explain both serial and parallel processing by adopting different parameter regimes. We present statistical methods to distinguish between serial and parallel processing based on both maximum likelihood estimates and decoding the momentary focus of attention when two stimuli are presented simultaneously. Results show that both processing mechanisms are in play for the simultaneously recorded neurons, but neurons tend to follow parallel processing in the beginning after the onset of the stimulus pair, whereas they tend to serial processing later on. This could be explained by parallel processing being related to sensory bottom-up signals or feedforward processing, which typically occur in the beginning after stimulus onset, whereas top-down signals related to cognitive modulatory influences guiding attentional effects in recurrent feedback connections occur after a small delay, and is related to serial processing, where all processing capacities are being directed towards the attended object.

**Author summary:** A fundamental question concerning processing of visual objects in our brain is how a population of cortical cells respond when presented with more than a single object in their receptive fields. Is one object processed at a time (serial processing), or are all objects processed simultaneously (parallel processing)? Inferring the dynamics of attentional states in simultaneously recorded spike trains from sensory neurons while being exposed to a pair of visual stimuli is key to advance our understanding of visual cognition. We propose novel statistical models and measures to quantify and follow the time evolution of the visual cognition processes right after stimulus onset. We find that in the beginning processing appears to be predominantly parallel, which develops into serial processing 150 – 200 *ms* after stimulus onset in prefrontal cortex of rhesus monkeys.

## 1 Introduction

A fundamental question in theories of visual search is whether the process is serial or parallel for given types of stimulus material (for comprehensive reviews, see [1–3]). In serial search, only one stimulus is attended at a time, whereas in parallel search, several stimuli are attended at the same time. The question of serial versus parallel search has been extensively investigated by behavioral methods in cognitive psychology, but it is still highly controversial. In this article, we briefly review extant empirical methods and their results and then present and exemplify a new method for distinguishing between serial and parallel visual search. The method is based on analysis of spike trains measured in prefrontal cortex of rhesus monkeys while being exposed to a pair of stimuli, which the animal should detect and later respond to with a saccade towards a target object, first presented in [4]. A spike train is a sequence of recorded times at which a neuron fires an action potential, and it is believed that spike times are the primary way to transmit information in the nervous system. Point process modelling is a natural mathematical framework for addressing such phenomena, and we embed this into models of visual attention, which provides means to quantify parallel versus serial visual processing.

### 1.1 Behavioral methods for distinguishing between serial and parallel visual search

In typical experiments on visual search, the task of the observer is to indicate as quickly as possible if a certain type of target is present in a display. Positive (target present) and negative (target absent) mean response times are analyzed as functions of the display set size *N* (the number of items in the display). The method of analysis was laid out by [5–7] and further developed by [8]. The foundation is as follows.

In a simple serial model, the *N* items are scanned one at a time. When an item is scanned, it is classified as a target or a distractor. The order in which items are scanned is independent of their status as targets versus distractors. A negative response is made when all items have been scanned and classified as distractors. Thus, the number of items processed before a negative response is made equals *N*. Furthermore, the rate of increase in mean negative response time as a function of *N* equals the mean time taken to process one item, Δ*t*. A positive response is made as soon as a target is found. Because the order in which items are scanned is independent of their status as targets or distractors, the number of items processed before a positive response is made varies at random between 1 and *N* with a mean of (1 + *N*)/2. Thus, the rate of increase in mean positive response time as a function of *N* equals Δ*t*/2 (see [9] for experimental evidence of serial processing in a behavioral task).

In a parallel model of attention, several stimuli can be attended at the same time. The first detailed parallel model of visual processing of multi-element displays was the independent channels model proposed by Eriksen and his colleagues (e.g., [10,11]). It was based on the assumption that display items presented to separated foveal areas are processed in parallel and independently up to and including the stage of pattern recognition. The independent channels model has been used to account for effects of *N* on error rates. The linear relations between mean response time and *N* predicted by simple serial models are difficult to explain by parallel models with independent channels. However, the linear relations can be explained by parallel models with limited processing capacity [12,13].

A multi-feature whole-report paradigm for investigating serial versus parallel processing was introduced in [14]: Suppose two features must be processed from each of two stimuli (i.e., a total of four features). Let processing be interrupted before all of the four features have completed processing. If, and only if, processing is parallel, there will be cases in which just one feature from each of the two stimuli completes processing before the interruption. This event, in which the observer has only partially encoded each of the two stimuli, should never happen when processing is serial. Thus, states with partial information from more than one stimulus are strong indicators of parallel processing. In the experiment of [14] (see [15] for replications and extensions), observers were presented with brief exposures of pairs of colored letters and asked to report both the color and the identity of each letter. The results showed strong evidence of states of partial information from each of the two stimuli (e.g., information of just the identity of one of the letters and just the color of the other one), and the results were fitted strikingly well by a simple parallel-processing model assuming mutually independent processing of the four features.

### 1.2 Method based on neurobiological data

As exemplified above, previous methods for distinguishing between serial and parallel visual search have been based on behavioral data, and the evidence obtained by these methods has been somewhat indirect. Moreover, these methods are all based on the assumption that processing is either serial or parallel, and that it stays the same throughout the trial. In this article, we present a new method for distinguishing between serial and parallel visual search, a method based on analysis of electrophysiological data. Furthermore, we propose measures to quantify the processing mechanism in a continuum between serial and parallel processing. The method relies on the probability-mixing model for single neuron processing [16], derived from the Neural Theory of Visual Attention [1,17], which states that when presented with a plurality of stimuli a neuron only responds to one stimulus at any given time. By probabilistic modeling and statistical inference using multiple simultaneously recorded spike trains, we infer and decode what the recorded neurons are responding to, providing a way to distinguish between parallel processing and serial processing on a neuronal level. The new method appears more direct than previous methods, and it is possible to analyze the time evolution of the processing mechanism over the course of a trial.

Consider an experiment in which we record the action potentials or spikes from each of a number of neurons of the same type, e.g., a set of functionally similar neurons in visual cortex with overlapping receptive fields (see, e.g., [16]), or neurons in prefrontal cortex that are believed to be dynamically allocated to process task-relevant information (see, e.g., [4]), which are the neurons we analyze in this paper. Suppose two stimuli (Stimulus 1 and 2) are both within the classical receptive fields of all of the recorded neurons, but otherwise the receptive fields are empty. In this situation, we may test whether processing is parallel in the sense that on any given trial, some of the recorded neurons represent Stimulus 1 throughout the trial, while others represent Stimulus 2 throughout the trial. We may assume that a neuron represents Stimulus 1 rather than Stimulus 2 if the likelihood of the observed spike trains becomes higher by assuming that the neuron represents Stimulus 1. We may also test whether processing is strictly serial (i.e., one stimulus at a time) by testing, for example, whether there is a time interval Δ_1_ in which all of the neurons represent Stimulus 1 and a time interval Δ_2_, nonoverlapping with Δ_1_, in which all of the neurons represent Stimulus 2.

Strictly parallel or strictly serial processing of two or more stimuli may hardly be expected in a biological system, and must be regarded as idealizations. However, we will show how to measure the goodness of approximation of search processes in the brain to simple serial and parallel search, as well as the time evolution of the processing mechanism.

In the following sections, we first present the statistical methods and probabilistic models that we employ to distinguish between parallel and serial processing, and then show and discuss the results obtained using both simulated data and experimental data.

In section 2, we introduce the experimental data that we rely on to measure the degree of parallel and serial processing in a realistic biological situation, and explain our proposed statistical criteria and measures to distinguish between parallel and serial processing. We propose two models, the hidden Markov model (HMM) and the correlated binomial model (CBM), to account for the spike train data in an attention framework, and calculate their likelihood functions. The maximum likelihood estimates (MLEs) provide a prior measurement of parallel versus serial processing. We then describe methods to decode the momentary focus of attention given the fitted models, which provides a posterior measurement. In section 3 we present the results of the analysis conducted on the experimental data.

## 2 Materials and Methods

We present two models that relate the theories of visual attention to neuronal behavior, providing a tool to distinguish or quantify between parallel and serial processing through spike train analysis. Under the assumption of serial processing, the neurons are correlated, acting together as a population. This dependence can arise through two different pathways: 1) There exists an underlying variable driving the neurons towards attending to the same stimulus, creating a dependence, even if the neurons are conditionally independent given the state of this underlying variable. 2) The neurons are directly positively correlated, driving them to synchronize their attention.

The first pathway is naturally described by a HMM, where the hidden Markov chain switches between different states influencing the neuronal attention. If time is discretized and there are two stimuli, this leads to a mixture of binomials at each discretized time step, where the number of components in the mixture distribution equals the number of states of the Markov chain. The binomial distributions provide probabilities of the number of neurons attending to each stimulus, in dependence of the hidden state of the Markov chain. The second pathway can be represented by a CBM, a mixture of an ordinary binomial and a modified Bernoulli [18], which is used independently at each discretized time step. For both models, the attended stimulus for each neuron is unobserved, and the inference is based on spike train data. We estimate parameters using MLE by marginalizing out the unobserved attention variables. The estimated parameters in either model describe neuronal properties and are used to obtain a *prior* measurement of the degree of parallel or serial processing. For both models, we also decode the hidden states from the posterior probabilities of the latent attention variables, i.e., an estimate of the stimulus the neurons were most probably attending to given their observed spike trains. The decoding of attentional behavior provides a *posterior* measurement of the degree of parallel or serial processing. The diagram in Fig 1 summarizes the flow of the analysis including parameter estimation, decoding and interpretation.

**Fig 1.**
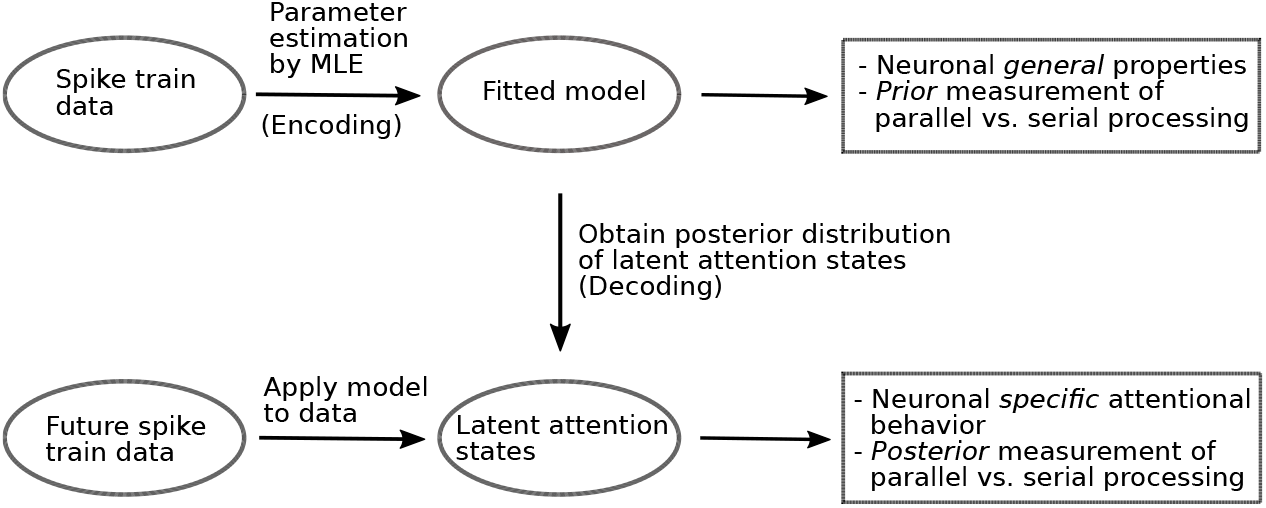
Flow diagram of the analysis.

### 2.1 Experimental Data

To distinguish between parallel and serial processing, we use the neural spike train data recorded from neurons in prefrontal cortex of two rhesus monkeys presented with two visual stimuli from [4]. They studied dynamic attentional construction, and found that in the early stage after stimulus onset when processing competing stimuli, the global attention is distributed among all objects with each neuron having a tendency towards its contralateral hemifield. In the late stage, the global attention is reallocated and neurons are redirected to the target stimulus. The data contain multiple simultaneously recorded neurons responding to two competing stimuli. The data are organized in daily sessions, and each session consists of a different set of recorded neurons. We only analyze the sessions where at least five neurons are recorded to have enough data to distinguish between parallel and serial processing, yielding a total of 48 sessions. The monkey fixed attention on a central red dot on a computer screen, then each trial began with a central cue indicating the target object of the specific trial. Each of two cues was paired with one of the two alternative targets. After a brief delay, a choice display was presented for 500ms containing two objects, one to the right and one to the left of the fixation point. The objects could be either the cued target (T), an inconsistent non-target (NI) because it was used as a target on other trials, a consistent non-target (NC) never serving as a target, or nothing but a gray dot (NO). After a brief delay, the monkey was rewarded with a drop of liquid for a saccade to the T location if a T had been shown, or if no T had been presented, for maintaing fixation (no-go response) for later reward. In the following we call a combination of two stimuli a *condition*. Table 1 shows the 12 possible conditions. The stimulus locations were denoted by whether they were contra-or ipsilateral with respect to the recorded neuron. S1 Fig a shows an example of the structure of the data in one session. To get an overall idea of the sample sizes, histograms in S1 Fig b and c show the average number of trials per condition over the 48 sessions, and the average number of simultaneously recorded neurons per trial over sessions, respectively.

**Table 1.** The 12 conditions used in the trials (combinations of stimuli).

| Condition | 1 | 2 | 3 | 4 | 5 | 6 | 7 | 8 | 9 | 10 | 11 | 12 |
| --- | --- | --- | --- | --- | --- | --- | --- | --- | --- | --- | --- | --- |
| con | T | T | T | NI | NC | NO | NI | NI | NC | NC | NO | NO |
| ipsi | NI | NC | NO | T | T | T | NC | NO | NO | NI | NI | NC |

Conditions can be merged into three groups, indicated by table cells: target in the contralateral side, target in the ipsilateral side, and all combinations with no target. Contra- and ipsilateral sides are with respect to the recorded neuron. T: Target; NI: Inconsistent non-target; NC: Consistent non-target; NO: No display.

We analyze the choice phase where the two stimuli are shown. In Fig 2 are shown the recorded spike trains of an example cell during this phase and 100 *ms* before and after. The red curves are kernel smoothing estimates of firing rates over time, plotted on top of the spike trains. The 12 subplots show the 12 conditions with the titles indicating stimulus on the contra- (left) and ipsilateral (right) sides with respect to the recorded neuron. The neuron seems to favor the target T with a higher firing rate, and its attention starts from the contralateral stimulus and is later redirected to the target stimulus, following the overall tendency of most neurons reported by [4]. In S2 Fig we show a complementary example neuron, which shows a tendency to the ipsilateral stimulus in the early stage, and later the attention is redirected to the target stimulus.

**Fig 2.**
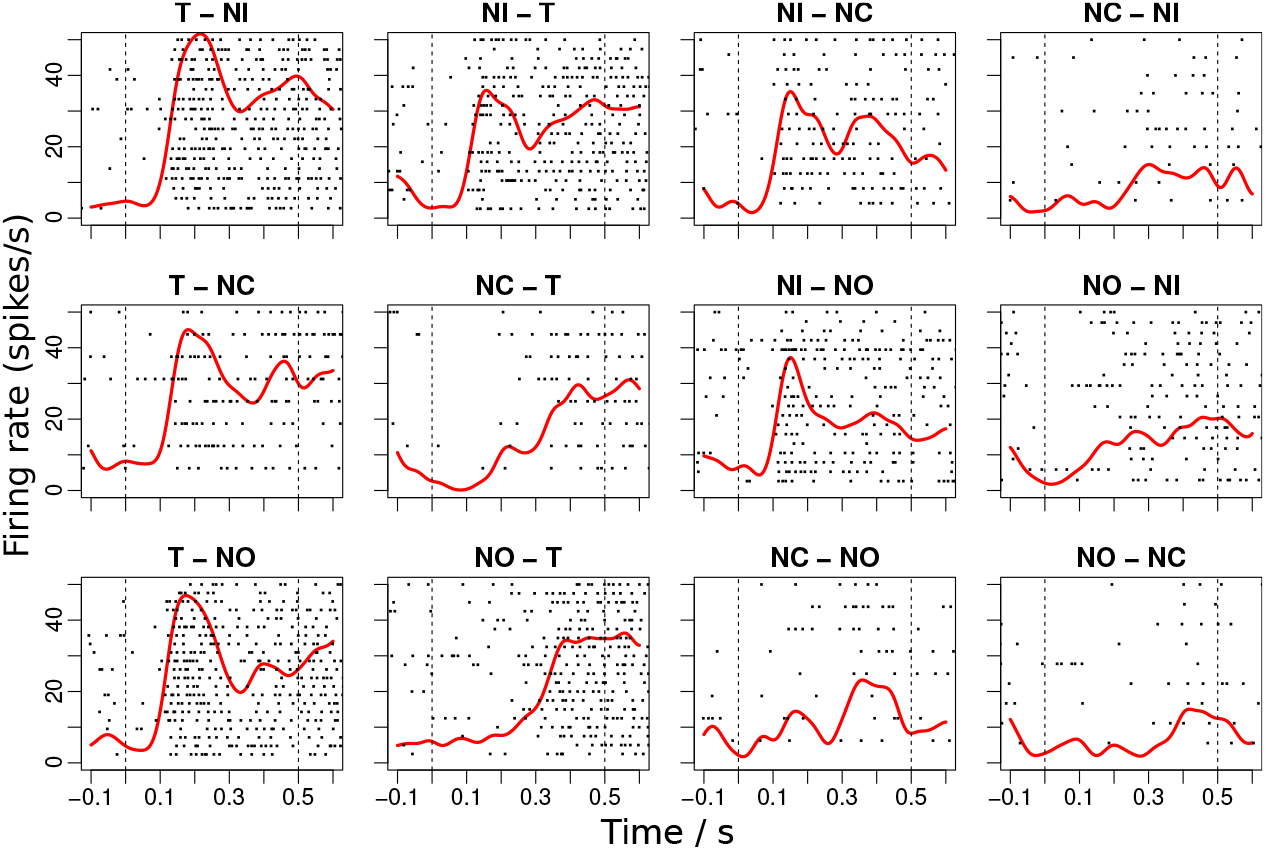
Raster plots of measured spike trains recorded from an example cell (MN1104lltask_3_0). The 12 conditions are indicated in the title of the subplot. Kernel smoothing estimates of the firing rates are shown in red. The stimulus in the left of the title indicates the stimulus of the contralateral side, and the right indicates the stimulus on the ipsilateral side with respect to the recorded neuron. The dashed lines indicate the interval of the choice phase where two stimuli are shown.

Furthermore, for this neuron there is more variability between trials under the same condition.

These figures present repeated trials of a single neuron. In S3 Fig, we show simultaneously recorded neurons in two conditions of an example session. The comparison of serial and parallel processing catches the difference among simultaneously recorded neurons within one trial in terms of their attended stimulus, which is hard or impossible to analyze by traditional methods by averaging across neurons and trials. We thus develop a new methodology modeling each single spike train and the correlation between spike trains. The serial and parallel processing can be distinguished using the estimated parameters. To account for neuronal response times, we discard the first 100 *ms* after stimulus onset, using the interval from 100 to 500 *ms* in the choice phase when estimating the parameters of the two models.

### 2.2 Measures for parallel versus serial processing

Here we define different measures of the degree of serial and parallel processing based on the estimated parameters of the models when a population of *n* neurons are presented with two non-overlapping stimuli in their receptive fields. These measures will vary with time, i.e., depend on the time since stimulus onset, but for ease of notation, we suppress time from the notation here. Later we will introduce the time dependency. We assume a homogeneous situation where all neurons follow the same distribution and are exchangeable, except for individual firing rates as responses to single stimuli. These measures are based on the basic probability-mixing model for the attention of single neurons employed in [16], where a neuron responds to a stimulus mixture with certain probabilities, such that the single neuron at any given time represents only one of the stimuli in the mixture. First, we consider the marginal distribution of the attended stimulus for each neuron. Let *p* denote the marginal probability of attending to one of the stimuli, say stimulus 1, such that the probability of attending stimulus 2 is 1 − *p*, where 0 < *p* < 1. If the neurons are independent, then the probability that all neurons attend the same stimulus is *p^n^* + (1 − *p*)^*n*^, and if the neurons are positively correlated, this is a lower bound of the probability that all neurons attend the same stimulus. Thus, *p* provides a measure of the tendency of serial or parallel processing. A narrow distribution (extreme probability, *p* either close to 0 or 1) favors serial processing, since in this case most neurons will attend the same stimulus. A wide distribution (non-extreme probability, p close to 0.5) favors parallel processing, since in this case neuronal attention will tend to split between the two stimuli. Second, we consider correlations between neurons. Since the neurons are exchangeable, the correlation coefficient, denoted by *ρ*, between any two neurons (pairwise correlation) is identical. Stronger positive correlation implies more tendency to serial processing, no matter what *p* is. Thus, if either the correlation is strong (*ρ* close to 1) or *p* is close to 0 or 1, serial processing is favored, while if both the correlation is weak and the probability is not extreme, parallel processing is favored. We summarize the different cases in Table 2.

**Table 2.** Effects of neural attentional probability and correlation to serial and parallel processing.

|  | Extreme probability | Non-extreme probability |
| --- | --- | --- |
| **Strong correlation** | Serial | Serial |
| **Weak correlation** | Serial | Parallel |

Extreme probability implies a probability close to 0 or 1, and strong correlation implies a correlation close to 1.

We now propose a single statistic as an alternative measure to distinguish between serial and parallel processing. Again, we suppose to have a stimulus mixture of two components and a population of *n* neurons attending to the mixture. The number of neurons, *Z*, attending to the first stimulus follows a distribution with probability mass function (PMF) *f*(*z*) for *z* ∈ {0,1,…,*n*}, such that *P*(*Z* = *z*) = *f*(*z*), which depends on the specific model. A distribution centered around *n*/2 indicates apparent parallel processing, and a distribution centered at 0 and/or *n* indicates apparent serial processing. Note that this population distribution incorporates both the marginal probability of attention of the single neurons and the correlation between neuron pairs. We define a statistic *D_n_* as a measure of the degree of serial or parallel processing, given by

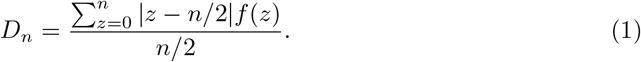

The statistic *D_n_* can be explained as a normalized expected deviation between the number of neurons attending to one stimulus and the half of the total number of neurons. If we split the neuron population according to which stimulus they attend giving two proportions (summing to 1), then *D_n_* is the average difference between the two proportions, and it can take values between 0 and 1. The smaller *D_n_* is, the more parallel processing is favored. The *D_n_* statistic depends on the total number of recorded neurons *n*. However, if we consider specific models for the PMF, for example the binomial models introduced below, the dependence of *n* can be removed by using the asymptotic version

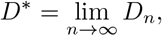

which provides a measure for the entire neuronal population relevant for the given task, not only the measured ones.

To summarize, to measure the degree of serial and parallel processing, we can use the attentional probability *p*, the correlation of neuronal attention *ρ*, and the deviation statistics *D_n_* or *D*\*.

### 2.3 Models

In this section we present two models to explain the spike train data in an attention framework. We discretize the 400 *ms* of the trial where both stimuli are presented, and which we use for the analysis, into *T* smaller intervals and let the models evolve dynamically over these intervals. Within any of these small time intervals, we assume that the attention of each neuron is not changing. Within a trial, let 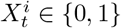 denote the attended stimulus of neuron *i* at time *t* for *i* = 1,…, *n*, *t* = 1,…,*T*, and let 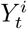 denote the spike train of neuron *i* in the *t*’th interval. We set 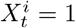 when neuron *i* attends stimulus 1 at time *t*, and 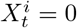 when attending stimulus 2.

#### 2.3.1 Hidden Markov Model and a Mixture of Binomial Distributions

To combine the visual attention hypotheses with neuronal dynamics, we adopt a HMM. The HMM assumes some underlying unobserved variable that drives the attention of the neurons. The HMM is defined over the *T* time steps. We let the probabilities *p* of the single neurons, which can be interpreted as attentional weights, depend on the state of the underlying HMM, which introduces correlation between neurons, even if they are conditionally independent given the hidden state, and the probabilities evolve over time following the dynamics of the HMM. Note that this implies that within each of the *T* intervals, model parameters governing the stochastic neuronal activity (the spike train generation) are constant. We use three hidden states, which can describe three attentional regimes: attention directed to the contralateral side, to the ipsilateral side, or equal attention to both sides. A transition between hidden states introduces a weight reassignment of the attention to the stimuli, and thus, new laws for the generation of spike trains. Let *C_t_* ∈ {1, 2, 3} denote the hidden state at time *t*. Fig 3 shows a diagram of the HMM for *T* = 3. Conditional on *C_t_*, the 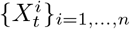 are independent.

**Fig 3.**
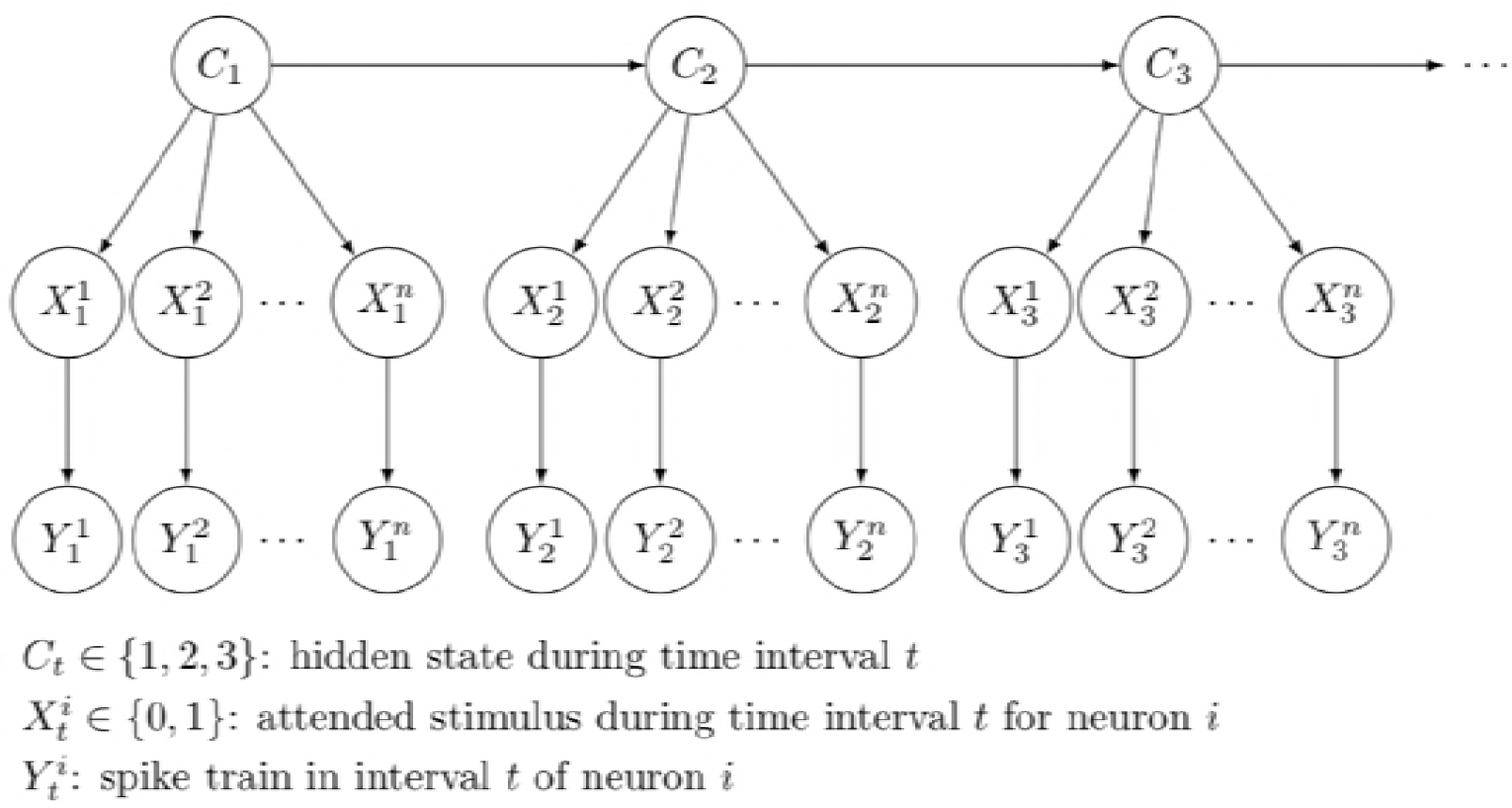
Diagram of the Hidden Markov Model. The HMM and attentional states from a group of *n* neurons to a mixture of two stimuli, using *T* = 3 discretized time steps.

Let the initial distribution of the Markov chain be given by **λ** and the transition probability matrix (TPM) by **Γ**:

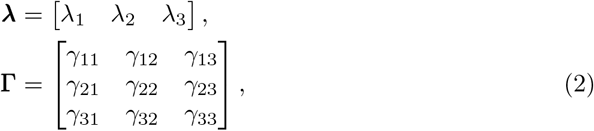

where 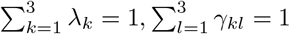 for *k* = 1, 2, 3, and λ_*k*_, *γ_kl_* ≥ 0 for *k, l* = 1, 2, 3. Here, *λ_k_* = *P*(*C*_1_ = *k*) and *γ_kl_* = *P*(*C*_*t*+1_ = *l*|*C_t_* = *k*) for all *t*. Let the vector *π_t_* = **λΓ**^*t*−1^ denote the distribution of *C_t_*, thus, *π_t,k_* = *P*(*C_t_* = *k*). The TPM **Γ** depends on the stimulus pair, but the initial distribution **λ** is only related to the location of the attended stimulus, since this is the initiation of the processing mechanism before the specific stimuli are perceived, and is thus the same for all stimulus pairs. However, we relax this assumption in Section 3.2.4. We denote by **Γ**_*m*_ the TPM of condition *m*.

Conditional on *C_t_*, neurons are assumed independent. Denote the probability of attending to stimulus 1 given state *c* by 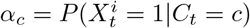, yielding the matrix:

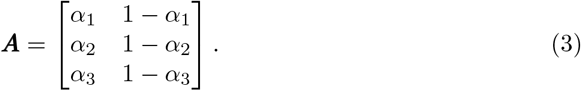

##### Attention probabilities and correlations

Calculating the probability distribution of *X_t_* is straightforward following the HMM. The vector

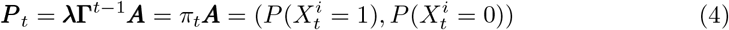

gives the vector of probabilities of attention to either stimulus. Straightforward calculations yield the moments and the correlation *ρ_t_* between 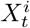 and 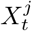:

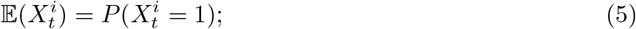

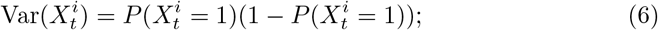

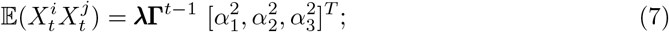

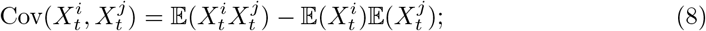

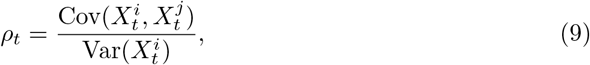

where ^*T*^ denotes transposition. The values 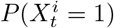 and *ρ_t_* can be used to measure the degree of serial and parallel processing as indicated in Table 2.

##### A mixture of three binomials

By marginalizing out the hidden state *C_t_*, the HMM structure implies that at each time point *t* the neuronal attention behavior for the *n* neurons follows a mixture of three binomial distributions, Bin3(*π_t_, α, n*). Here, *α* = (*α*_1_, *α*_2_, *α*_3_) are the probability parameters of the three binomials, and the weights are given by *π_t_*. The number of binomial trials equals the number of simultaneously recorded neurons *n*. The PMF for the mixture of three binomials is

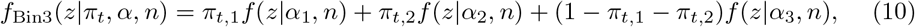

where 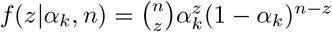 is the PMF of the binomial distribution.

The *D_n_* statistic is calculated using Eq (1). For the mixture of three binomials in (10), the asymptotic version is given by

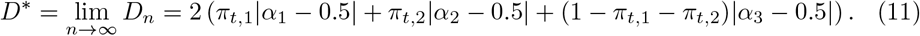

Fig 4a illustrates how the probability and the correlation affect serial and parallel processing for the HMM using *n* =10 neurons for four different parameter settings. The parameter settings are shown in Table 3, together with the corresponding calculated attention probabilities, correlations, *D*_10_ and *D*\* values. Only when *p* is not close to 0 or 1 and the correlation is weak, the 10 neurons tend to split between the two stimuli, indicating parallel processing. Otherwise, a majority of the neurons attend to the same stimulus, suggesting serial processing. The four cases from Case 1 to 4 show increasing degree of parallel processing. In all cases are *D*\* < *D_n_*, so if more neurons are involved, we expect more clear parallel processing for the given parameters.

**Fig 4.**
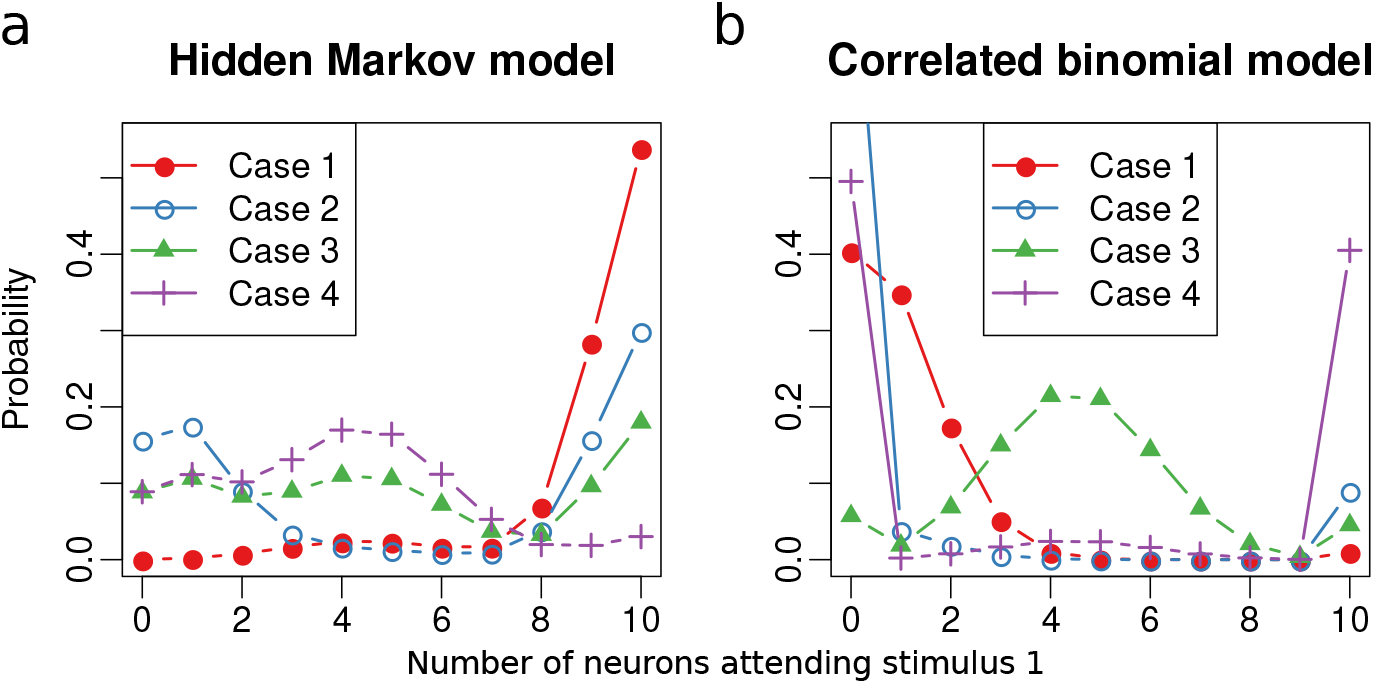
The probability mass function of the number of neurons attending to stimulus 1. The four sets of parameter values are given in Table 3. a) HMM leading to a mixture of three binomials. b) CBM.

**Table 3.** Parameter values, probabilities of attention, correlation, and the deviation statistics *D*_10_ and *D*\* for the HMM and the CBM.

| | $\pi_{t,1}$ | $\pi_{t,2}$ | $\pi_{t,3}$ | $P(X_t^i = 1)$ | $\rho$ | $D_{10}$ | $D^*$ |
| --- | --- | --- | --- | --- | --- | --- | --- |
| Hidden Markov model, $\alpha = (0.95, 0.45, 0.1)$ | | | | | | | |
| Case 1 | 0.9 | 0.1 | 0.0 | 0.9 | 0.25 | 0.836 | 0.82 |
| Case 2 | 0.5 | 0.05 | 0.45 | 0.543 | 0.691 | 0.823 | 0.815 |
| Case 3 | 0.3 | 0.45 | 0.25 | 0.513 | 0.407 | 0.586 | 0.515 |
| Case 4 | 0.05 | 0.7 | 0.25 | 0.388 | 0.165 | 0.426 | 0.315 |
| Correlated binomial model. |  |  |  |  |  |  |  |
| Case 1 | - | - | - | 0.1 | 0.1 | 0.82 | 0.82 |
| Case 2 | - | - | - | 0.1 | 0.9 | 0.98 | 0.98 |
| Case 3 | - | - | - | 0.45 | 0.1 | 0.33 | 0.19 |
| Case 4 | - | - | - | 0.45 | 0.9 | 0.93 | 0.91 |

#### 2.3.2 Correlated Binomial model

In the CBM the neurons are assumed directly correlated. It was studied in [18,19], and is denoted by CBin(*n,p, ρ*), where *n* is the number of correlated Bernoulli trials (simultaneously recorded neurons in our model setting), 0 < *p* < 1 is the probability 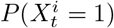, and *ρ* is the correlation coefficient. In this model the number of neurons *z* attending stimulus 1 follows a mixture of two distributions. One is an ordinary binomial distribution with parameters *n* and *p*. The other is a fully correlated distribution where *z* ∈ {0, *n*}, which can be viewed as a modified Bernoulli distribution with support {0, *n*} with parameter *p*. The weight of the Bernoulli component is the correlation coefficient *ρ*. The probability mass function is given by

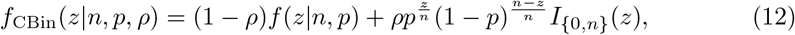

where *I*_{0,*n*_}(*z*) is the indicator function which equals 1 for *z* ∈ {0, *n*} and 0 otherwise.

As before, we assume the distribution at the first step identical for all stimulus pairs, whereas at all later steps, the distribution depends on the stimulus pair. Thus, at *t* = 1 the simultaneously recorded neurons follow CBin(*n, p*_1_, *ρ*_1_), and at *t* > 1 they follow CBin(*n, p_t,m_, ρ_t,m_*) for stimulus pair *m*. We do not assume a dependence structure over time, as in the HMM, and the behavior at each time step is independent of the behavior at other time steps. Instead, the correlation between simultaneously recorded neurons are modeled directly by the parameter *ρ*. Compared with the HMM, where the correlation is described through the attentional reassignment with a Markov chain, the CBM is more direct.

Denote by *C_t_* the hidden index, indicating either the binomial (*C_t_* = 1) or the Bernoulli (*C_t_* = 2) component in the mixture. The probability of attention is directly obtained from the parameter *p_t,m_*, and the correlation is obtained from *ρ_t,m_*. The asymptotic version of the deviation statistic *D*\* to measure the degree of serial and parallel processing is given by

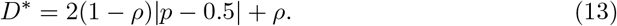

Fig 4b shows the PMF of the correlated binomial distribution for four different parameter settings, shown in Table 3, together with the *D*_10_ and *D*\* values.

### 2.4 Likelihood functions and model fitting

The spike trains are modelled by point processes using conditional intensity functions (CIF) [20,21], see also [16]. Suppose a spike train y in the interval [*T_s_,T_e_*] contains the spike times *y* = {*t*_1_, *t*_2_,…} with *T_s_* ≤ *t*_1_ < *t*_2_ < ⋯ ≤ *T_e_*, and that it attends to the same stimulus during the entire interval. The probability of observing *y* given the attended stimulus *x_t_* is given by [21, 22]

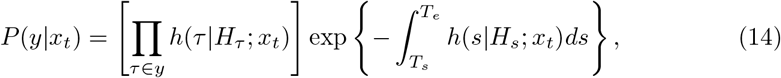

where *H_s_* is the spike history up to time *s*, and *h*(*s*|*H_s_; x_t_*) is the conditional intensity function, which we model using

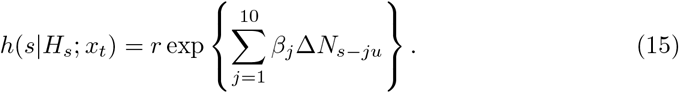

The base firing rate *r* := *r^i^* is neuron specific and a function of the attended stimulus and the location (contra-or ipsilateral). Note that only the attended stimulus is relevant, not the condition. For each neuron, there are therefore 7 rate parameters, representing T, NI and NC at either side, and a parameter for NO. The exponential term models the influence of the past spikes during the previous 10ms on the neuronal activity. The constant *u* =1 *ms* is the discretization unit determined by the experiment, and Δ*N_t_* denotes whether there is a spike (Δ*N_t_* = 1) or not (Δ*N_t_* = 0) in the time interval [*t,t* + *u*). For simplicity, we assume that only past spikes of the neuron itself have an effect. All neurons are assumed to share the same set of *β* parameters *β_j,j_* = 1, 2,…, 10.

Let 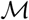 denote the considered conditions (stimulus pairs) and let 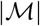 denote the number of conditions. For simplicity, we do not always distinguish between all 12 conditions shown in Table 1, but sometimes merge them into classes, such that there will be fewer parameters to estimate. In particular, we will consider the three classes of conditions indicated in the table, defined by whether there is a target in the stimulus pair, and if there is, whether it is contra-or ipsilateral. Under condition *m*, let the set 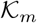 contain all the conducted trials. In trial *k*, let the set 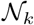 contain all the simultaneously recorded neurons and let 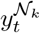 denote the spike trains from these neurons in the *t*’th interval, and likewise for the hidden attentional states 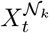. Each 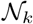 is a subset of the set of all neurons 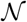 used in the session, 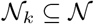, because not all neurons are used in all trials (S1 FigA).

#### 2.4.1 Hidden Markov Model

We denote the conditional probability of the 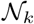 spike trains at time *t* given *C_t_* by a diagonal matrix:

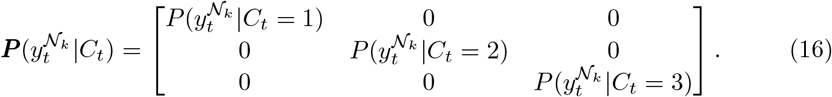

The likelihood function of all spike trains in one session is then given by

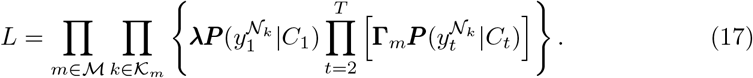

By conditioning on the hidden attentional states 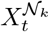, we obtain

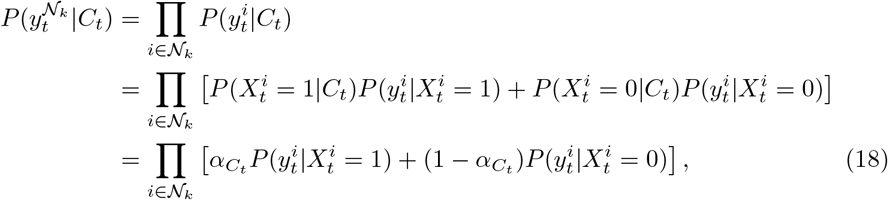

where 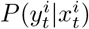 is given in Eq. (14). We obtain MLEs of the parameters by maximizing the likelihood function. The parameters to be inferred are summarized in Table 4.

**Table 4.** Parameters to be estimated for each session in the HMM and the CB models.

| Name | Explanation | Dimension |
| --- | --- | --- |
| **Hidden Markov Model** |  |  |
| $\lambda_m = [\lambda_1 \quad \lambda_2 \quad 1 - \lambda_1 - \lambda_2]$ | Initial distribution, the same for all conditions $\mathcal{M}$ | 2 |
| $\Gamma_m = \begin{bmatrix} \gamma_{11}^m & \gamma_{12}^m & 1 - \gamma_{11}^m - \gamma_{12}^m \\ \gamma_{21}^m & \gamma_{22}^m & 1 - \gamma_{21}^m - \gamma_{22}^m \\ \gamma_{31}^m & \gamma_{32}^m & 1 - \gamma_{31}^m - \gamma_{32}^m \end{bmatrix}$ | Transition probability matrix for each condition $m \in \mathcal{M}$ | $6\vert \mathcal{M}\vert $ |
| $\mathbf{A} = \begin{bmatrix} \alpha_1 & 1 - \alpha_1 \\ \alpha_2 & 1 - \alpha_2 \\ \alpha_3 & 1 - \alpha_3 \end{bmatrix}$ | Conditional probability of neuronal attention | 3 |
| $r^i$ | Base firing rates, different for each neuron in $\mathcal{N}$ | $7\vert \mathcal{N}\vert $ |
| $\beta$ | Weights in the CIF model, the same for all neurons $\mathcal{N}$ | 10 |
| **Correlated Binomial Model** |  |  |
| $\rho_{t,m}$ | Correlation coefficients for $m \in \mathcal{M}$ and $t = 1, \dots, T$ | $\vert \mathcal{M}\vert \cdot (T - 1) + 1$ |
| $p_{t,m}$ | Probability parameter for $m \in \mathcal{M}$ and $t = 1, \dots, T$ | $\vert \mathcal{M}\vert \cdot (T - 1) + 1$ |
| $r^i$ | Base firing rates, one for each neuron in $\mathcal{N}$ | $7\vert \mathcal{N}\vert $ |
| $\beta$ | Weights in the CIF model, the same for all neurons $\mathcal{N}$ | 10 |

#### 2.4.2 Correlated binomial model

Under the CBM, the attention of the simultaneously recorded neurons follow a mixture of a binomial and a modified Bernoulli. The likelihood of the spike trains in condition *m* at time *t* in trial *k*, 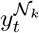, is given by

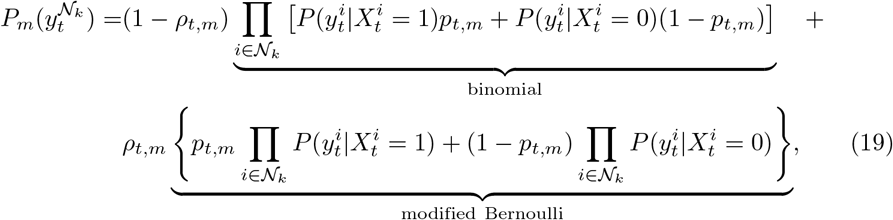

where 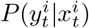 is given in Eq. (14). The likelihood of the data of an entire session is

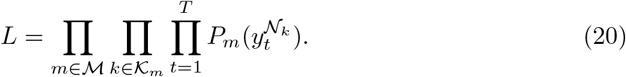

The parameters are summarized in Table 4.

We summarize the differences of the HMM and the CBM in Table 5. In both models, it is assumed that in the early stage, i.e., the first discretized interval from 100 *ms* to 100 + 400/*T ms*, neuronal attention is only affected by the position of stimuli (ipsi-or contralateral) and not by stimulus types (T, NI, NC or NO). This assumption is supported by the empirical findings by firing rate averaging showing attentional reallocation over time [4]. It is also assumed that under the same stimulus types, the attentional parameters are identical, implying that in all the trials of one condition, neurons follow the same distribution, and differences from trial to trial are due to randomness.

**Table 5.** Differences between the Hidden Markov model and the correlated binomial model.

|  | HMM | Correlated binomial |
| --- | --- | --- |
| Motivation | Extends the probability-mixing model with dynamic weight reassignment. | Treats neuronal attention as correlated binomial variables. |
| Neuronal correlation | Described through the Markov chain. | Modeled directly by parameters. |
| Parameter dimension | $13 + 2\vert \mathcal{M}\vert + 7\vert \mathcal{N}\vert $ | $12 + 2\vert \mathcal{M}\vert (T - 1) + 7\vert \mathcal{N}\vert $ |
| Meaning of $C$ | Hidden state of the Markov chain, each state giving different stimulus weights. Implies top-down control of attention. | State of neurons being either completely independent or fully positively correlated. |

### 2.5 Decoding the attentional state

Decoding means to infer the attended stimulus from the observations and the estimated parameters. To show the main idea, we suppress time and neuron indicator from the notation for the moment, denoting the hidden state by *C*, the attended stimulus by *X* and the spike train data by *Y*. The posterior of *X* given *Y* = *y* is

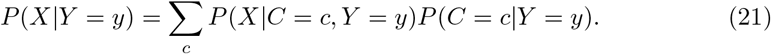

The strategy is to first estimate *P*(*C* = *c*|*Y* = *y*) and then *P*(*X*|*C* = *c, Y* = *y*) conditional on *C* = *c*. We are particularly interested in the PMF and the deviation statistic of the attended stimuli, which we can calculate using *P*(*X|C* = *c, Y* = *y*) for different states *C*. In the following, the decoding is explained for the two models in more detail.

#### Decoding in the Hidden Markov model

First we decode the hidden states *C_t_* in the HMM model. It is performed at each discretized time step by the forward-backward algorithm. Let 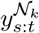 denote the spike trains in intervals *s* to *t*, for 1 ≤ *s* < *t* ≤ *T* in trial *k*, where 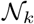 denotes the simultaneous recorded neurons in the *k*’th trial. The probability of *C_t_* conditional on the observed spike trains at all time intervals 1 to *T* can be expressed as

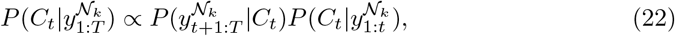

where

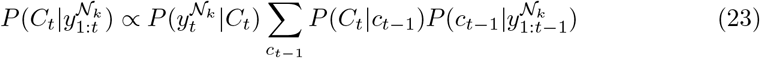

is the forward probability, calculated recursively by a forward sweep over 1 to *T*, and

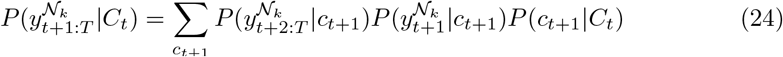

is the backward probability, calculated recursively by a backward sweep over 1 to *T*. When calculating the forward and backward probabilities, the likelihood conditional on the hidden state, 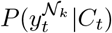, is obtained by conditioning on the neuronal attention 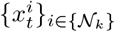:

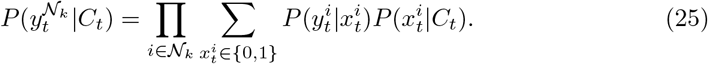

After decoding the hidden state 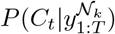, the next is to decode 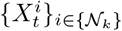 conditional on *C_t_*:

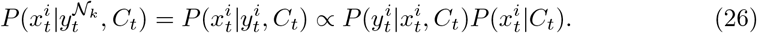

For all spike trains in trial *k*, 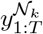, we have thus obtained the discrete posterior distributions of the hidden states 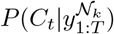 and the attended stimulus of each spike train 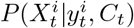, at all time steps *t* = 1,…, *T*. This yields the marginal posterior 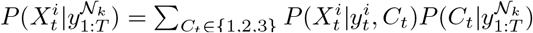.

At each time step *t*, conditional on *C_t_*, spike trains are independent and the posterior probabilities 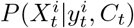 are different from spike train to spike train. Thus, the attended stimuli of all neurons follow a Poisson binomial distribution, a generalization of the ordinary binomial distribution where each Bernoulli trial has a distinct success probability [23]. The PMF of the Poisson binomial distribution is calculated numerically using methods from [24]. Marginalizing out *C_t_*, at each time step t we then have a mixture of three Poisson binomial distributions. The PMF of this mixture distribution can be regarded as probabilities of the number of neurons that have attended stimulus one, conditional on their observed spike trains. The deviation statistic *D_n_* can also be obtained from the PMF.

#### Decoding in the correlated binomial model

In the CBM, spike trains between different time steps and different trials are independent (except for the memory component, the exponential term in Eq. (15)). Thus, decoding can simply be done independently for each discretized time step in each trial. Now, let *C_t_* be an index indicating either the binomial or the Bernoulli component in the mixture. As previously, we first decode *C_t_* by calculating 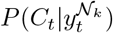, then find the PMF by calculating 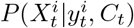. We have

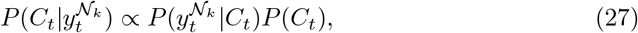

where the two cases *C_t_* = 1 and *C_t_* = 2 are given by the two components in Eq. (19). Then for each case of *C_t_* we decode the attended stimulus 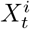. When *C_t_* = 1, i.e., the binomial case, 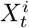 is obtained for each spike train independently with 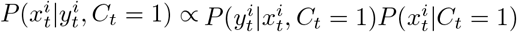, resulting in a Poisson binomial distribution. When *C_t_* = 2, i.e., the fully correlated Bernoulli case, the attended stimuli of all neurons are the same, which is obtained by 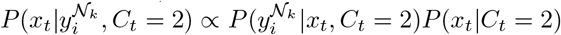, and the result is still a modified Bernoulli. Finally, the PMF is a mixture of a Poisson binomial and a modified Bernoulli.

## 3 Results

### 3.1 Simulated data

We first simulate spike train data and check if our models and methods work properly on the simulated data. For both the HMM and the CBM, we consider three parameter settings. In all cases, we use 10 simultaneously recorded neurons, repeated for 20 trials. The parameters, including base rates and response weights, are the same for the three cases. We consider only one stimulus condition, such that each neuron only has two base rate parameters, one for the contralateral and one for the ipsilateral sides.

The parameter values used in the simulations are shown in Table 6. For the HMM, we use a time step of 0.1s and a total of 10 time steps. For the CBM, we use a time step of 0.1s and a total of 5 time steps. S4 Fig shows the probabilities, correlations, *D*\* and *D_n_* values as functions of time. Simulated example spike trains are also shown.

**Table 6.**
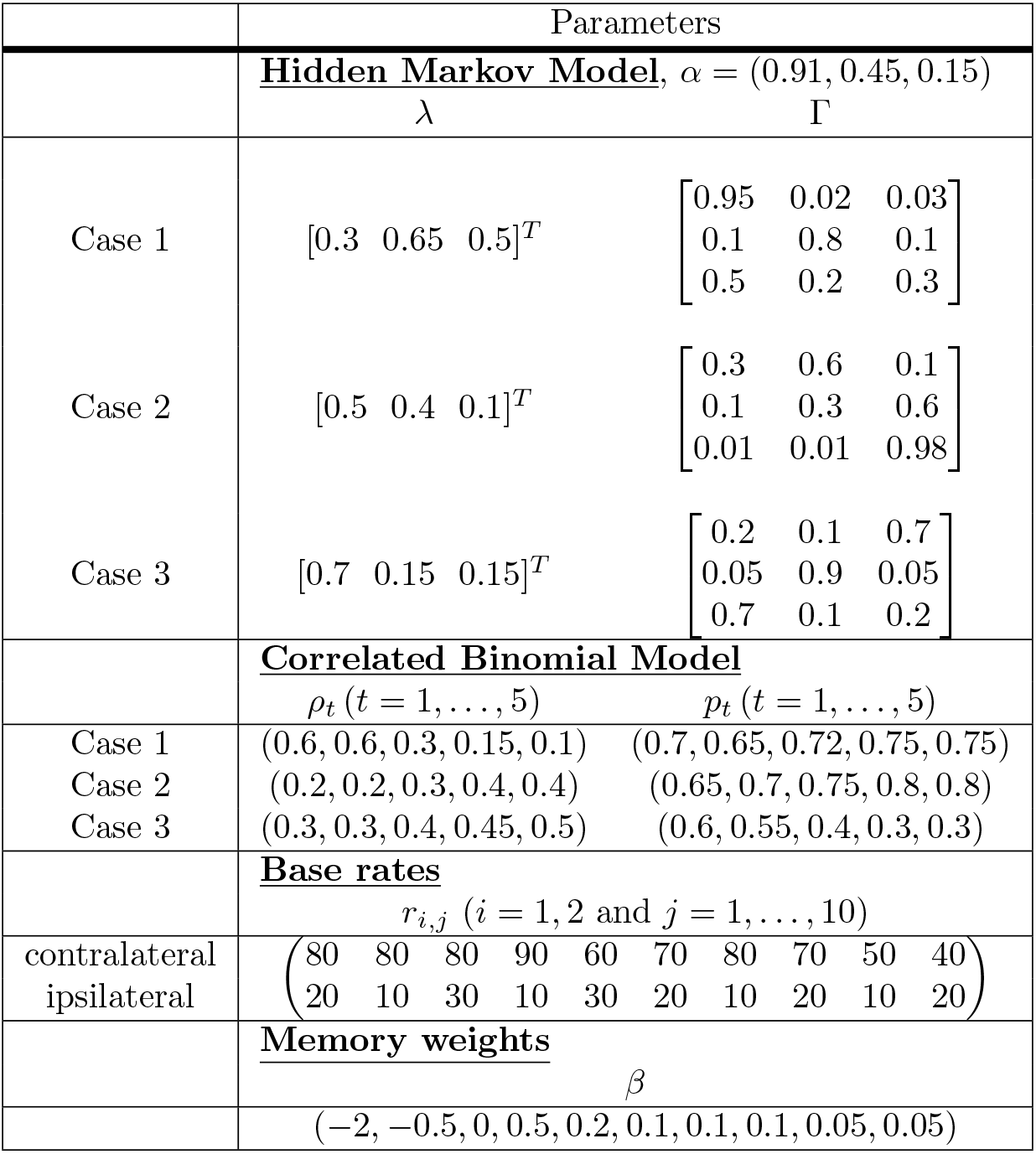
Parameters used to simulate data from the HMM and the CBM. Different *ρ* and *p* parameter values are used at different time steps *t*. Firing rates and weight values for the 10 simulated neurons are the same in the two models. The base firing rate is denoted by *r_i,j_* for stimulus *i* and neuron *j*. The contralateral stimulus is represented by *i* = 1 and the ipsilateral stimulus by *i* = 2. The memory weights are the same for all neurons, denoted by *β*.

We apply the model fitting to the simulated data, and the MLEs are shown in Figs 5 and 6 together with the true parameters for the HMM and the CBM. The simulation and model fitting procedure are repeated 100 times.

**Fig 5.**
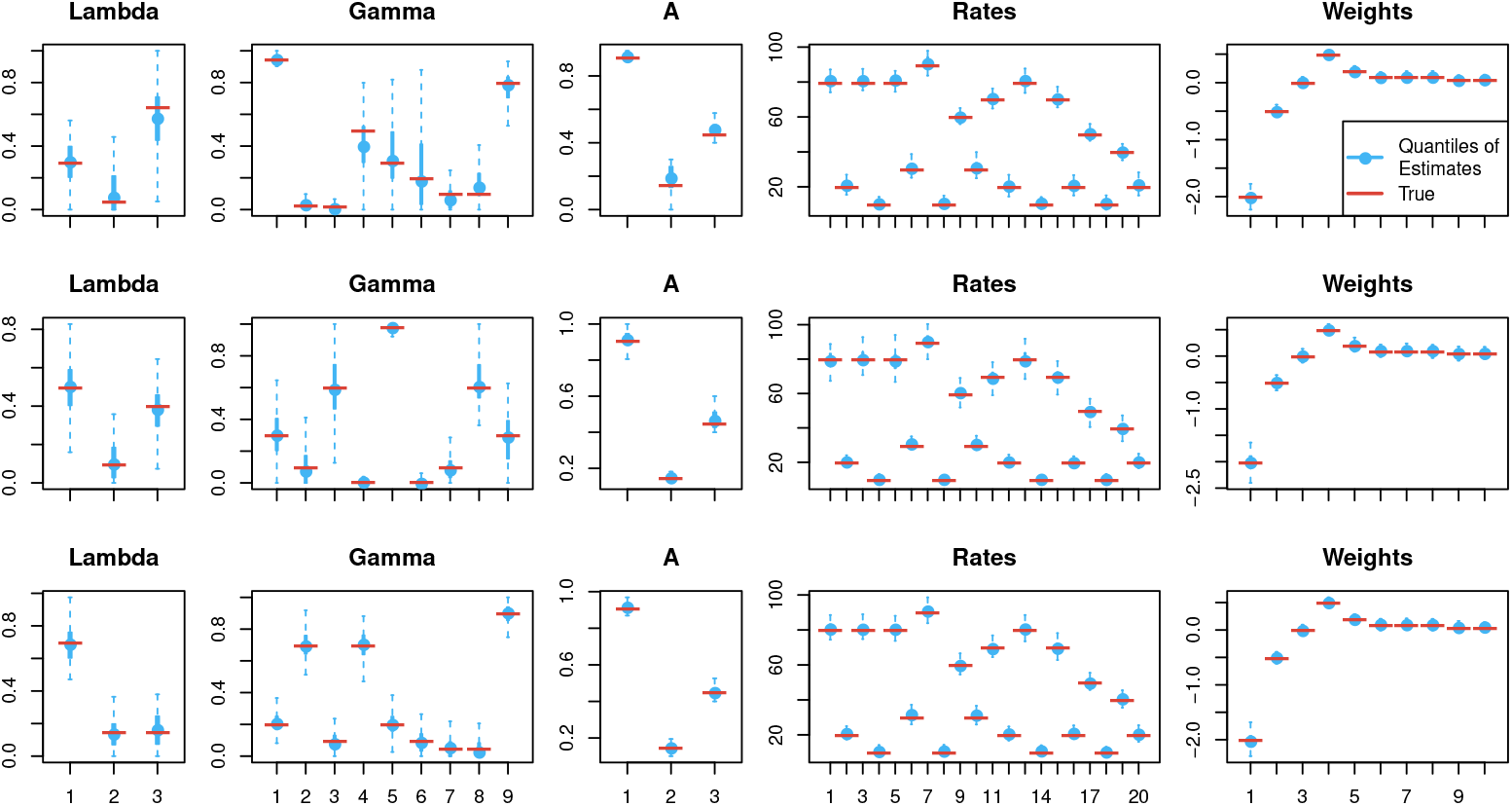
Parameter estimates of the hidden Markov model from the simulation study. The estimates are shown as quantiles of the 100 repetitions. The dashed lines represent the full 0 – 100% quantiles, and the solid lines represent the 25% – 75% quantiles. The central dot is the median. The red lines are the true values used in the simulation. The upper, middle and lower panels represent the three cases.

**Fig 6.**
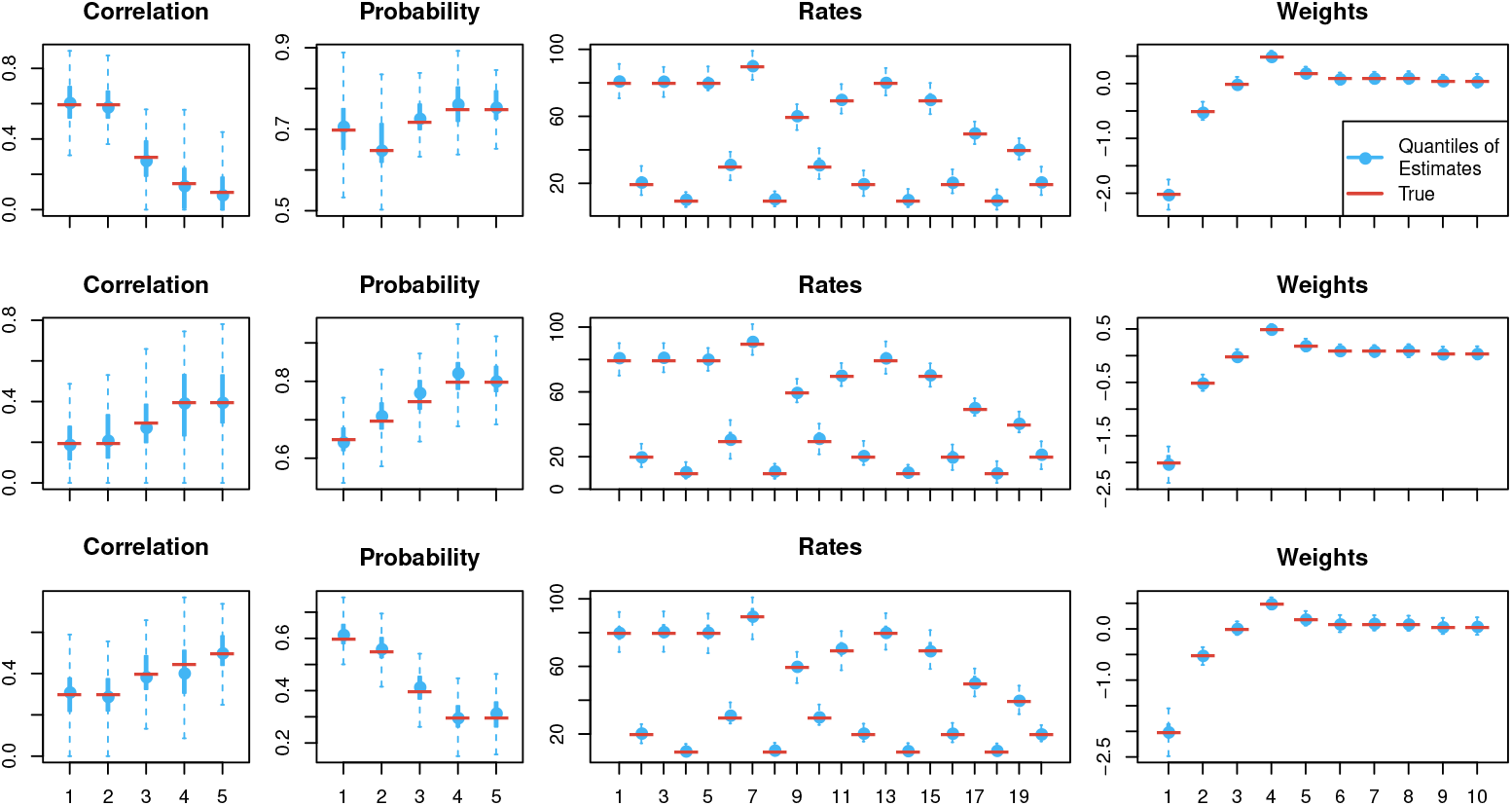
Parameter estimates of the correlated binomial model from the simulation study. The estimates are shown as quantiles of the 100 repetitions. The dashed lines represent the full 0 – 100% quantiles, and the solid lines represent the 25% – 75% quantiles. The central dot is the median. The red lines are the true values used in the simulation. The upper, middle and lower panels represent the three cases.

The *D*\* and *D_n_* values are computed from the parameter estimates, and are shown in Fig 7 together with the true *D*\* and *D_n_* values.

**Fig 7.**
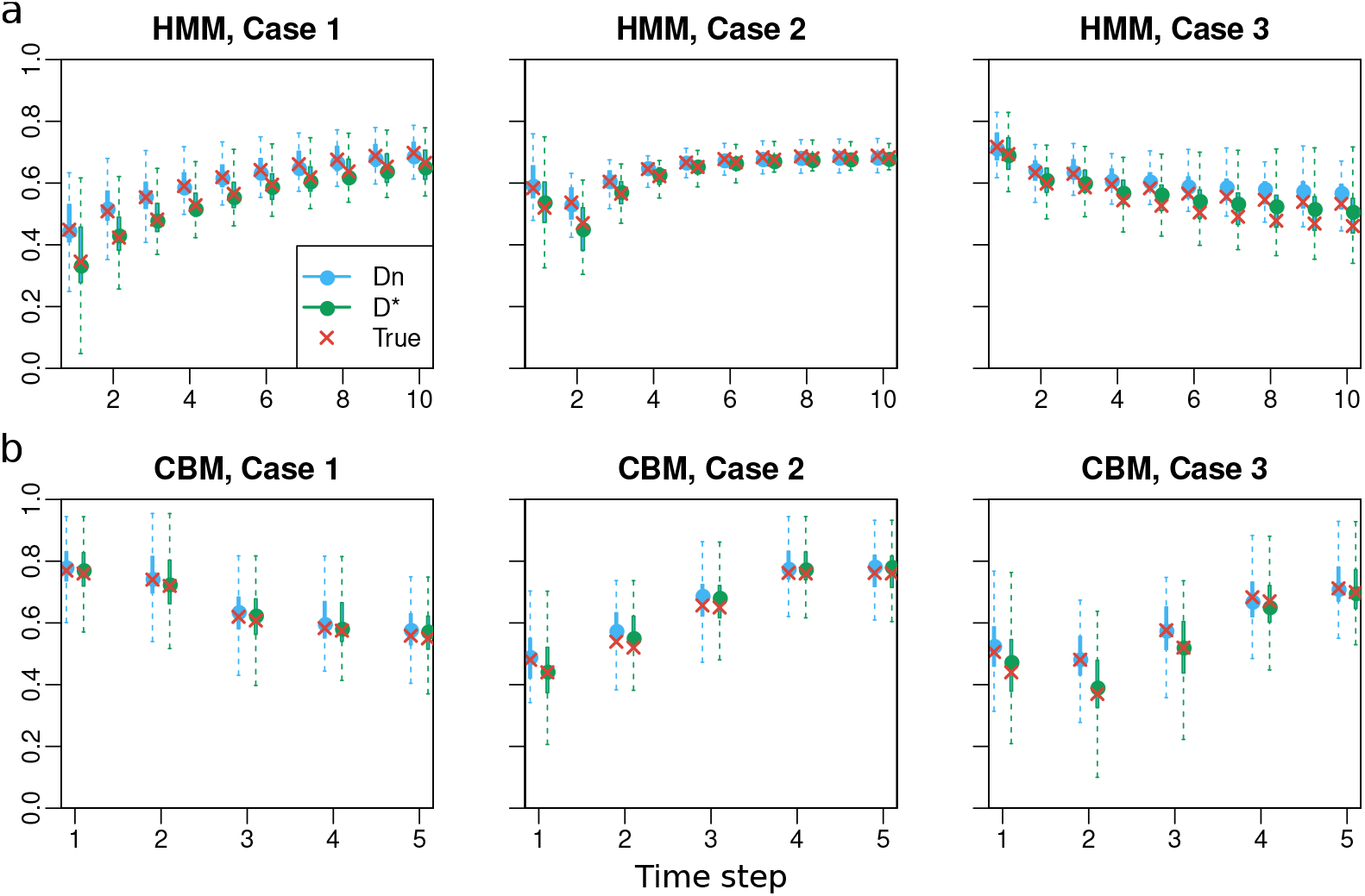
Deviation statistics values computed from parameter estimates and true parameters. The estimates of *D_n_* are shown in blue and the estimates of *D*\* are shown in green as quantiles of the 100 repetitions. The dashed lines represent the full 0 – 100% quantiles, and the solid lines represent the 25% – 75% quantiles. The blue dots are the medians. The red dots are the true values used in the simulation. a) HMM. b) CBM.

Finally, we also perform decoding analysis using the estimated parameters for each trial. The *D_n_* values from the decoding are plotted in Fig S5 Fig together with the *D_n_* values computed directly from the parameter estimates.

The conclusion from this simulation study is that parameters can be successfully estimated, and the *D_n_* and *D*\* values computed from the parameter estimates are close to the true values. The *D_n_* values from the decoding analysis have large variances, due to the small sample size of 10 neurons. However, the median *D_n_* values from the 100 decoding repetitions are often close to the encoding results.

### 3.2 Experimental data

The experimental spike train data from [4] were fitted to both models. For a discretization with *T* steps, an equal length of 400/*T ms* were assigned to all time intervals. Three different discretizations of *T* = 3, 5 or 10 were used, and two different classes of conditions with either all 12 or only 3 classes determined by whether there is a target in the stimulus pair, and in that case, whether it is contra-or ipsilateral (see Table 1). The models were fitted to each of the 48 sessions independently.

#### 3.2.1 Parameter estimation in HMM

Fig 8 illustrates parameter estimates for the HMM using different condition and step number settings. Fig 8a shows the probability of attending to the stimulus at the contralateral side, *p_t_* = *P*(*X_t_* = 1), as kernel density plots from all 48 estimates. Three line types (solid, dashed and dotted) indicate the three time steps, and four colors represent four types of conditions. At *t* =1 all conditions follow the same distribution, so there is a single black curve. For the subsequent time steps, the condition types are: stimulus pairs with T on the ipsilateral side; stimulus pairs with T on the contralateral side; stimulus pairs with NO on the ipsilateral side; and stimulus pairs with NO on the contralateral side. It illustrates that neuronal attention slightly prefers the contralateral stimulus in the beginning right after stimulus onset (the black density curve is centered slightly towards larger values than 0.5), and later on tends to follow T and avoid NO. Note that here we conduct model inference using all 12 conditions, and only combine similar conditions together for presentation.

**Fig 8.**
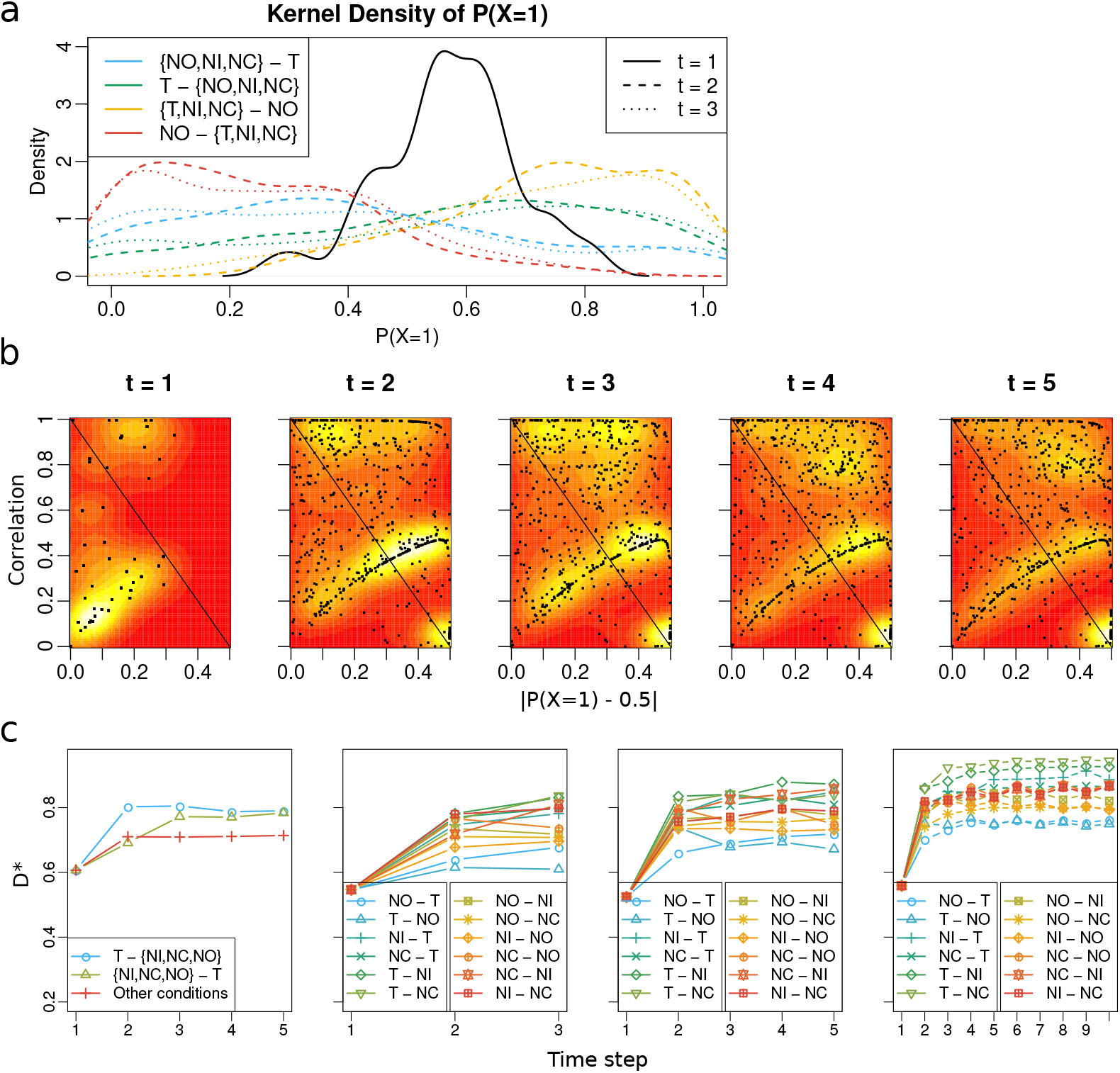
Experimental data: Results for the HMM. a) Kernel density representation of the estimates of *P*(*X* = 1), i.e., the probability of a neuron attending to the contralateral stimulus, obtained using *T* = 3 and all 12 conditions. b) Correlation estimates vs probability extremeness estimates at the different time steps, on top of a two-dimensional kernel density estimate as heatmaps, obtained using *T* = 5 and all 12 conditions. c) Estimates of *D*\* using 3 merged conditions with *T* = 5 (left), all 12 conditions with *T* = 3 (middle left), *T* = 5 (middle right), and *T* =10 (right).

In Fig 8b, the estimates of the correlation *ρ_t_* are plotted against the estimates |*p_t_* – 0.5| (difference of the probability of the contralateral stimulus from 0.5, or probability “extremeness”) for each time step *t* = 1,…,5, on top of a two-dimensional kernel density estimate (bandwidth: 0.25) of the points as heatmaps. There are 48 estimates in the leftmost panel at *t* = 1 (no difference between conditions), and 48 × 12 estimates in the remaining panels from 12 conditions in 48 sessions. A straight line is plotted on the anti-diagonal for easier reading. The lower left region of the heatmap represents a tendency of parallel processing, and all other regions represent a tendency of serial processing. In the leftmost panel corresponding to the first time step, a big portion of the estimates fall in the lower left region. At later time steps, the estimates tend to move to the right and upper regions. This implies that, in an early stage stimuli tend to be processed in parallel. Later on more and more neurons share the same attended stimulus in the form of serial processing. Despite the moving tendency, there are points lying on both sides of the straight line at all time steps. This is evidence supporting both processing mechanisms for all time steps throughout the entire spike train.

In Fig 8c we investigate the asymptotic deviation statistic *D*\*. The average *D*\* is calculated over the 48 session estimates for each condition. The left panel shows the *D*\* values obtained from parameter estimates using 3 classes of merged conditions with *T* = 5. The remaining panels show results using all 12 conditions, with *T* = 3, 5 and 10 for the middle left, the middle right and the right panels. In all cases, *D*\* grows larger over time, implying stronger serial processing. Further, different settings of discretization and condition merging give different results. The differences caused by using a larger *T* may be due to smaller sample sizes (shorter spike trains with only few spikes).

#### 3.2.2 Parameter estimation in correlated binomial

The estimates of the CBM is shown in Fig 9. The results are similar to the HMM. In Fig 9b, we see apparent parallel processing at *t* = 1, while later on the correlation for most estimates goes to either 1 or 0, and the probability becomes more extreme. For *t* > 1, in most of the 48 × 12 estimates the weight parameter of the mixture (the correlation coefficient) is close to either 1 or 0, meaning one component is dominating over the other. This is because of the small number of simultaneously recorded neurons in most trials (see S1 Fig), which is insufficient for obtaining good estimates in a mixture model. This is a weakness of the CBM since it only contains two extreme components representing either full independence or full correlation. Model fitting of the CBM on limited sample sizes can bias the correlation parameter. To check this suspicion, we looked at the estimates from session “MN110411”, the right-most neuron in S1 Fig *c* with the largest number of simultaneously recorded neurons, and found that the estimates of the correlation lie almost uniformly across 0 to 1, indicating that the estimates of either 0 or 1 of the correlation in other sessions can be an artefact of small sample sizes.

**Fig 9.**
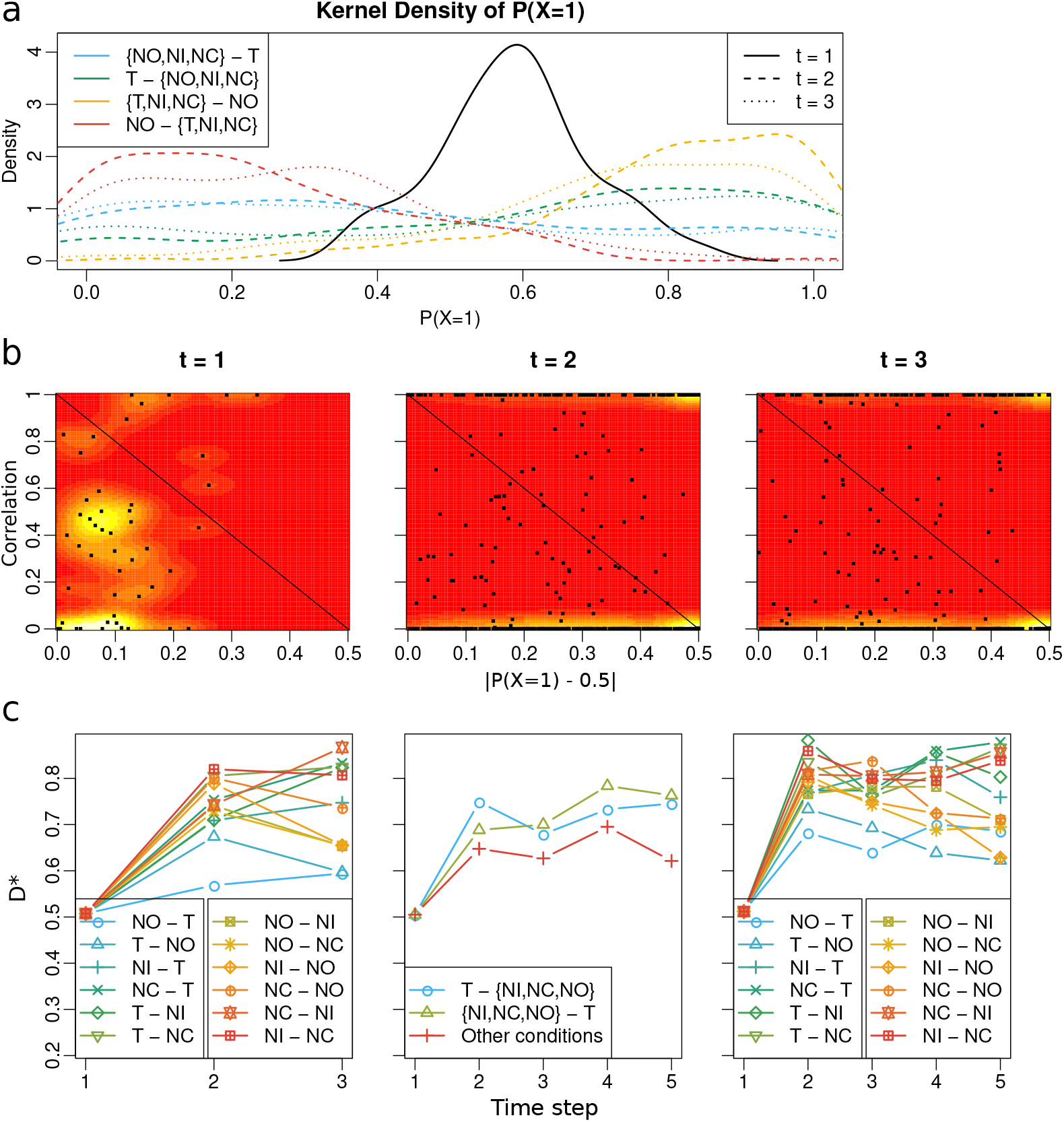
Experimental data: Results for the CBM. Fig a and b use all 12 conditions with *T* = 3. Fig c uses 12 conditions with *T* = 3 (left), 3 merged conditions with *T* = 5 (middle), and 12 conditions with *T* = 5 (right). See caption of Fig 8 for explanation.

#### 3.2.3 Decoding

Here we decode the attended stimulus of each neuron conditional on the observed spike trains. The parameters used in the decoding algorithms are the estimated parameters obtained by MLE. In the HMM model we show results using *T* = 3, *T* = 5 and *T* = 10, and in the CBM using only *T* = 3.

Fig 10 shows the decoding of the attended stimulus for an example trial containing 10 simultaneously recorded spike trains in session “MN110411”, condition NO-T. The same data set is decoded using the HMM with *T* = 3, 5 and 10, and the CBM with *T* = 3. The values of the *D_n_* statistics are calculated based on the decoded probabilities of a single trial containing simultaneously recorded neurons.

**Fig 10.**
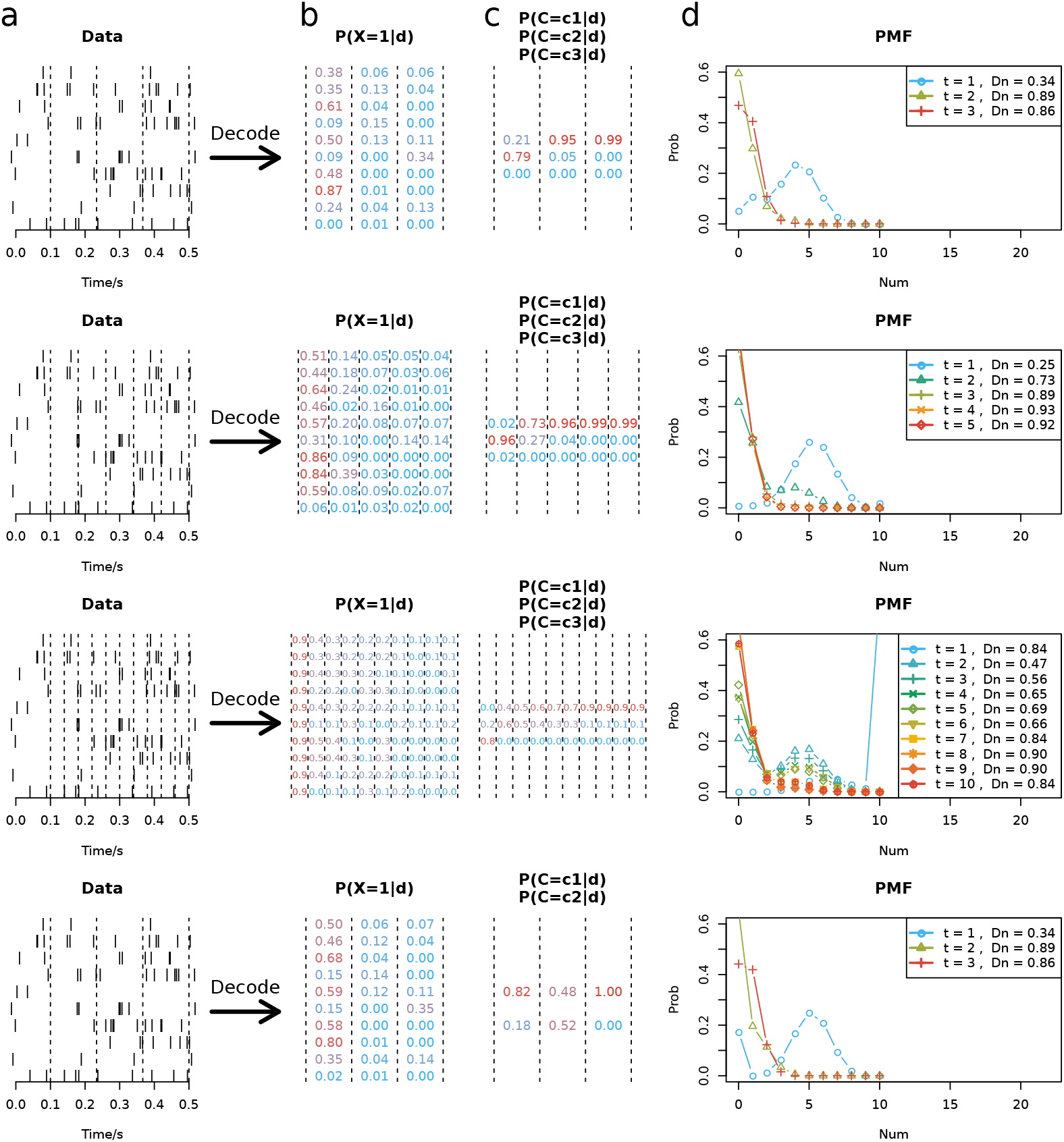
Decoding of an example trial using different models. All models use all 12 conditions. The top three panels show the results of the HMM model using *T* = 3, 5 and 10. The bottom panel shows the CBM with *T* = 3. a) Ten simultaneously recorded spike trains from a trial in session “MN110411”. The dashed lines indicate the discretization. b) The posterior probability of each spike train attending the contralateral stimulus at each time step, with the dashed lines indicating the time steps corresponding to the discretization in a. Estimates in red color indicate higher probability of attending the contralateral stimulus and blue color indicates higher probability of attending the ipsilateral stimulus. Note that the target is located in the ipsilateral side. c) The posterior probability of the hidden state *C* responsible for top-down control. For the HMM, the hidden state indicates the index of the binomial component, and for the CBM, the first hidden state is the independent binomial component and the second is the fully correlated Bernoulli component. d) The PMF of the number of neurons attending to the contralateral stimulus conditional on the spike train data for each time step, with the *D_n_* values shown in the legend. These values are calculated by Eq. (1) using the estimated PMFs.

In Fig 11 we show box-plots of the *D_n_* values from multiple trials across all sessions as a function of time. Again we include four cases: HMM with *T* = 3, 5 and 10, and CBM with *T* = 3. If a trial has too few simultaneously recorded spike trains, the *D_n_* values will be biased, and there will be large variance across trials. For example, in the simulation study in Fig 4, we see large quantiles for the decoding results even with very good parameter estimates. For this reason, we only consider trials with at least 10 simultaneously recorded spike trains. Note that the minimum number in a trial here is different from the number of simultaneously recorded neurons in a session, because in many trials not all simultaneously recorded neurons are used. We pre-selected data such that the number of simultaneously recorded neurons in a session is at least 5, but in most trials the simultaneously recorded spike trains can be fewer. We see a similar trend as in the encoding results: The *D_n_* values are increasing with time, indicating stronger serial processing. The degree of serial processing becomes maximal starting from around the middle of the spike train. In particular, for HMM with *T* = 5 (top right), the *D_n_* box-plot at *t* = 2 shows smaller median and larger variance than *t* = 3. For HMM with *T* = 10, the box-plots at *t* = 2, 3, 4 show large variance (reaching low values) compared with *t* ≥ 5. Note that the similar results in Figs 8c and 9c are *prior* measures based on estimated parameters, and the plots in Fig 11 are *posterior* measures based on the decoded attended stimulus for specific spike train data. Finally, in all models and at all time steps, there is evidence of both parallel and serial processing, implied by the wide box-plots.

**Fig 11.**
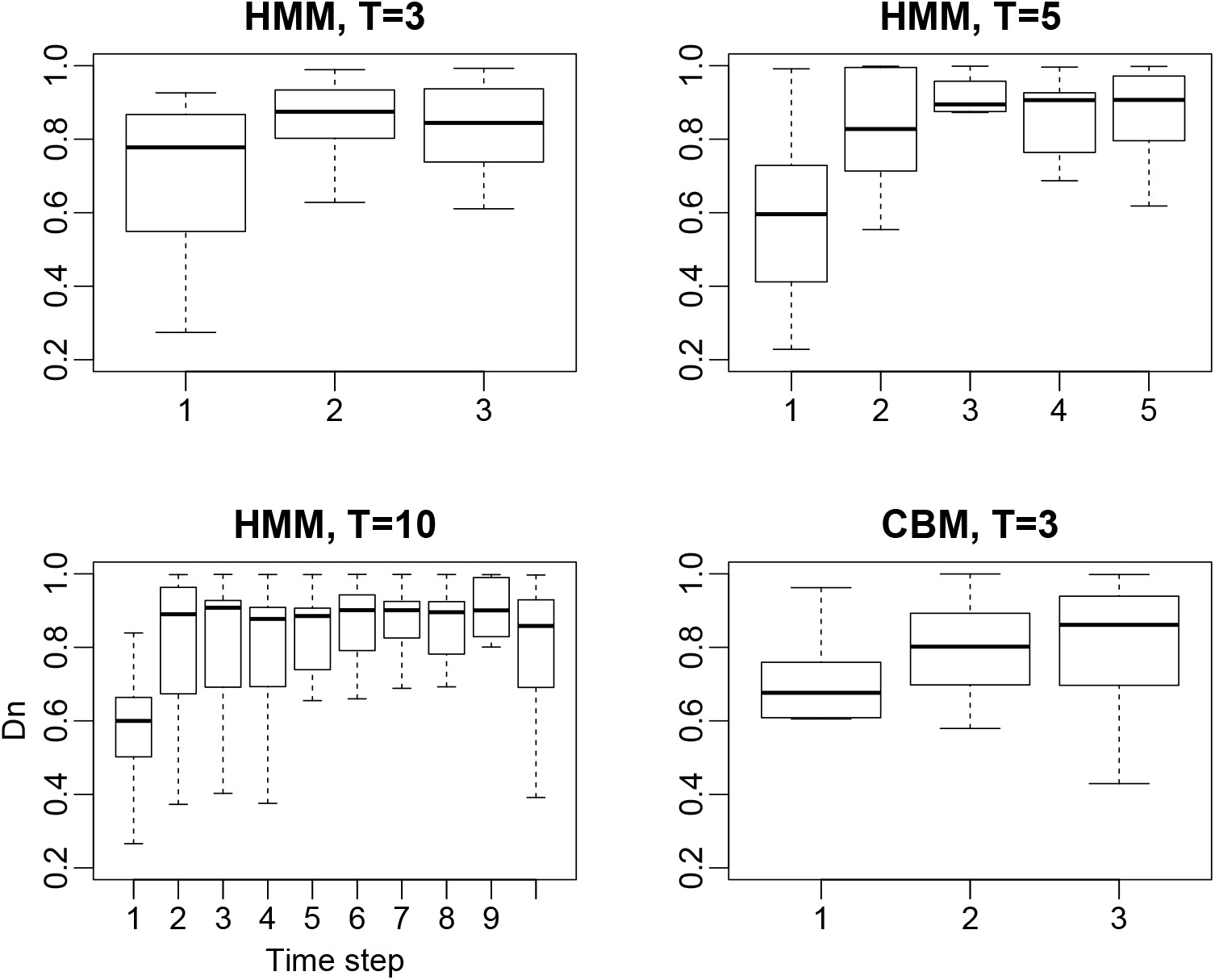
Decoding results of *D_n_*. Only trials with more than 10 simultaneously recorded neurons are included. Results are from the HMM using *T* = 3, 5 and 10, and the CBM using *T* = 3, respectively, indicated by the titles of the figures.

#### 3.2.4 Specific initial probabilities for each condition

Previously we have used the same initial probabilities for all conditions, i.e., neuronal attention in the beginning right after stimulus onset is only affected by stimulus locations and not by stimulus types, which is supported by the original study [4]. Here we conduct a further analysis discarding this assumption and allowing each condition to have its own initial probabilities. Doing so will greatly increase the number of parameters to estimate, and the estimation and inference results of the mixture models become increasingly unreliable given limited data size and large noise. We only analyze the most simple example for the HMM using three time steps with three merged conditions. In Fig S6 Fig are shown the *D*\* statistics obtained using parameter estimates, similar to Figs 8c and 9c, but for the two settings: fixing the same initial probabilities or assuming different initial probabilities for each condition. Though the *D*\* results are different in the two plots in Fig S6 Fig, the conclusion remains the same; neuronal attention is more parallel right after stimulus onset and becomes more serial later on.

## 4 Discussion

In this study we combine the point process neuron models describing spike trains with the neural interpretations of serial and parallel processing hypotheses in visual search. We propose a HMM and a CBM to describe neuronal attention in neurophysiological measurements from prefrontal cortex in rhesus monkeys. Results show that parallel processing is favored in some sessions while serial processing is favored in other sessions, and there is evidence for both parallel and serial processing at all time steps. Overall, we see a tendency towards parallel processing in the early stage after stimulus onset, and serial processing in the late stage. This means that, right after stimulus onset, neurons tend to split to attend different stimuli, and later neurons become more synchronized sharing the same attended stimulus. Furthermore, at the early stage neurons prefer the contralateral stimulus, while in the late stage neurons favor the T and avoid NO, which agrees with the study conducted by averaging across spike trains [4].

The early state of parallel processing can be related to feedforward or bottom-up processing, where the sensory inputs are being processed before higher level cognitive modulatory influences of recurrent feedback or top-down processing has begun [25,26]. In the later stage, where top-down signals have had time to modulate the attention, the neural activity tends to synchronize around the attended object, resembling serial processing. Similar results have been observed in event-related potentials in electroencephalography (EEG) measurements [27]. They found that forward connections are sufficient to explain the data in early periods after stimulus onset, whereas backward connections become essential after around 220 *ms*. Even if the exact timing of the switch between bottom-up and top-down signals is not clear, there is evidence that after 200 *ms* back projections play a prominent role, even if selective responses are elicited already after 100 *ms* after stimulus onset (see [26] and references therein). Quantification of the relative contribution of feedforward and feedback signals characterizing visual perception remains unclear, and thus, the concepts of parallel and serial processing and our suggested analysis tools provide a useful mean for elucidating these questions.

Decoding analysis provides posterior probabilities of neuronal attentions, yielding an estimate of the PMF and therefore also of *D_n_*. This can be used to analyze attentional behavior for any given simultaneously recorded spike trains in future trials. The conclusions regarding parallel and serial processing from the overall distribution of *D_n_* on all trials and sessions from the decoding analysis are the same as in the prior analysis using only parameter estimates. Note that although both the prior and posterior analysis provide similar results, the conclusions regarding neuronal attentional properties should be drawn from the prior analysis based on the MLE. The MLE gives the optimal estimation of the neuronal properties based on all the available data. The decoding analysis, on the other hand, estimates what the neuron’s attention could have been during a specific trial based on the data from this trial, and the uncertainty of the decoding is represented by posterior distributions.

In [4], parallel processing in the early stage was reported. The same conclusion is drawn from our analysis, where we find that the neurons prefer the contralateral stimulus in the early stage, and integrating both hemispheres gives simultaneous parallel processing. Furthermore, there exists not only such parallel processing considering the whole brain, but also parallel processing based on neurons in a single recording site, as supported by our finding. Though the simultaneously recorded neurons in one location show a tendency towards the contralateral stimulus in the early stage, there is strong evidence showing they split their attention between stimuli located on both sides in a parallel way.

The models here are fitted to the specific data set from [4] and the model structure contains the experimental conditions specific for this data set. However, with trivial adjustments, the models also apply to generic neurophysiological data that consist of simultaneously recorded spike trains. Currently the models and methods only support two stimuli, and a future extension is the generalization to an arbitrary number of stimuli.

The two models, the HMM and the CBM, yield different results regarding the degree of serial and parallel processing. This is partly because the two models are based on different assumptions. The biological reality of attention, which we try to describe with these simple models, is complicated, and the two models approximate the reality and explain neural attention from different perspectives. Further, the experimental data are noisy with limited sample size and the models contain a large number of parameters, which leads to large variance of estimators. For one trial or session, the difference between the two models could be large, but the overall results of the two models over a large number of sessions produce similar conclusions. However it makes more sense to make comparisons under the same model. For example, we compare different conditions or different time steps only under the same model.

Another issue is the variability between sessions for the same model. We assume the whole prefrontal area follow a probabilistic model and we want to estimate the model parameters. However, in each session we only have a small subset with 5 to 20 simultaneously recorded neurons from a recording site, and the number is even smaller for single trials (S1 Fig), with each neuron having its distinct firing rate and attentional pattern (Figs 2 and S2 Fig). Thus, there is a large variance of the estimates from session to session, and we obtain the overall result by averaging and applying kernel density estimation. To obtain more stable and accurate results it would be beneficial to use a larger simultaneously recorded population of neurons.

## Supporting information

**S1 Fig.**
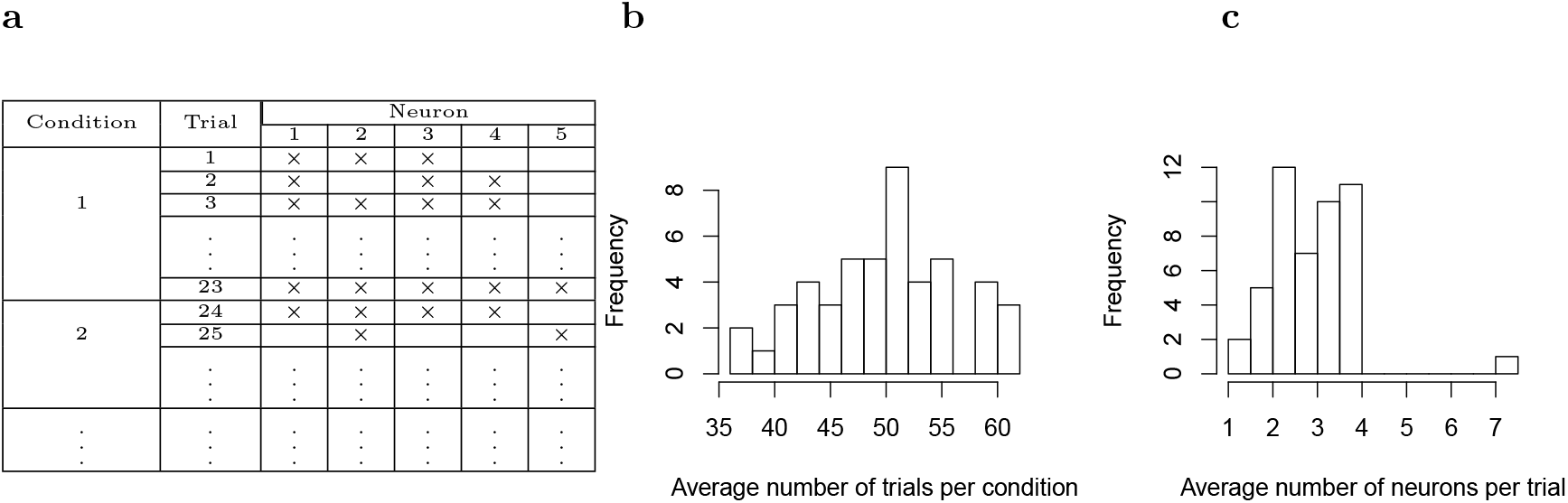
Data structure and sample sizes. a) Example of recorded neurons and condition within each trial in a daily session. The symbol × indicates that the neuron is recorded in the given trial. In this session, five neurons are recorded. Condition 1 was used in 23 trials, and in trial 1 and 2 three neurons are recorded, but not the same ones. b) Average number of trials per condition in 48 sessions. c) Average number of neurons per trial in 48 sessions. In all sessions, at least 5 neurons are recorded, however, not all enter in each trial. Histograms are based on 48 numbers (one for each session).

**S2 Fig.**
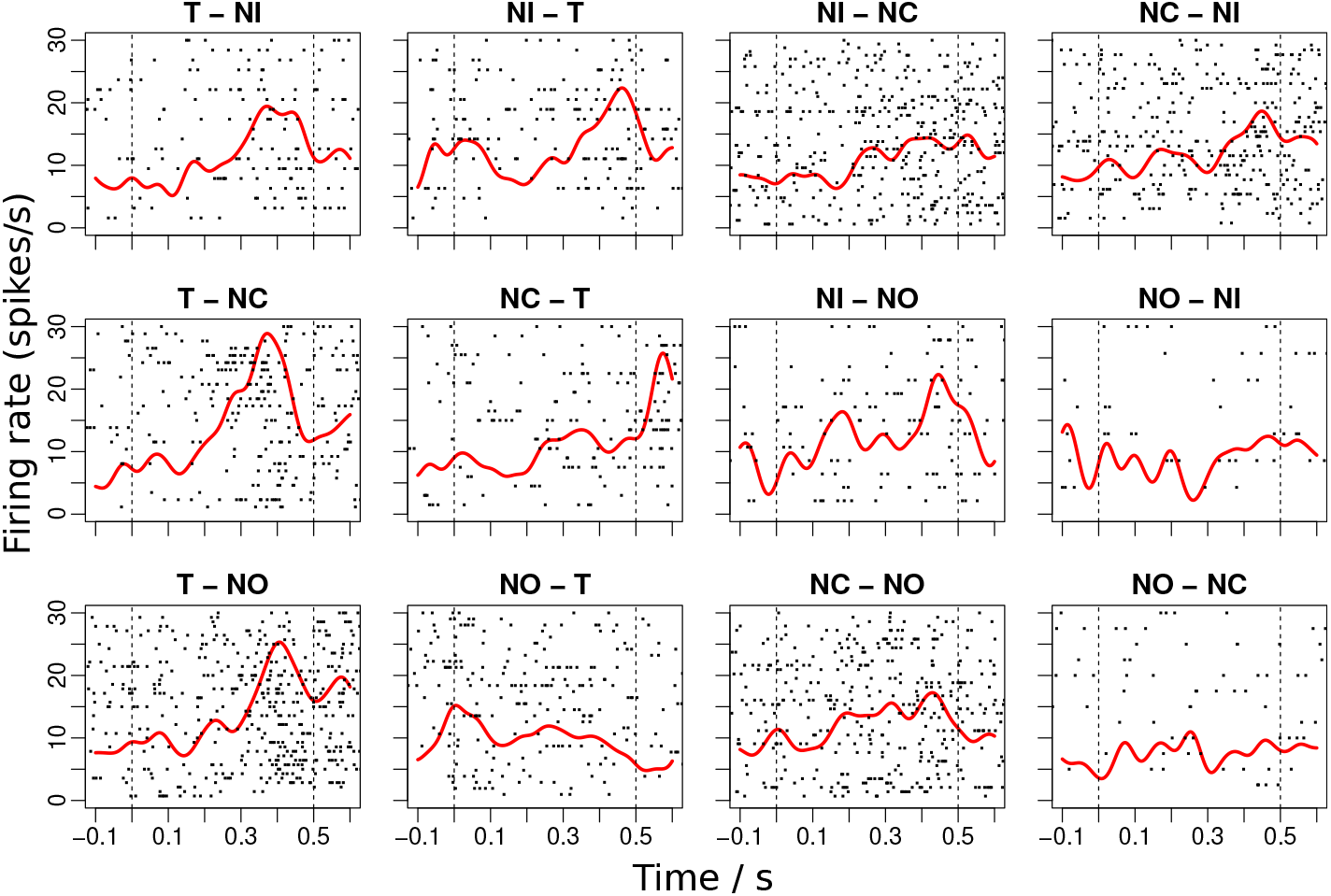
Raster plots of measured spike trains recorded from an example cell (mj081029a_8_0). The 12 conditions are indicated in the title of the subplot. Kernel smoothing estimates of the firing rates are shown in red. The stimulus in the left of the title indicates the stimulus of the contralateral side, and the right indicates the stimulus on the ipsilateral side with respect to the recorded neuron. The dashed lines indicate the interval of the choice phase where two stimuli are shown.

**S3 Fig.**
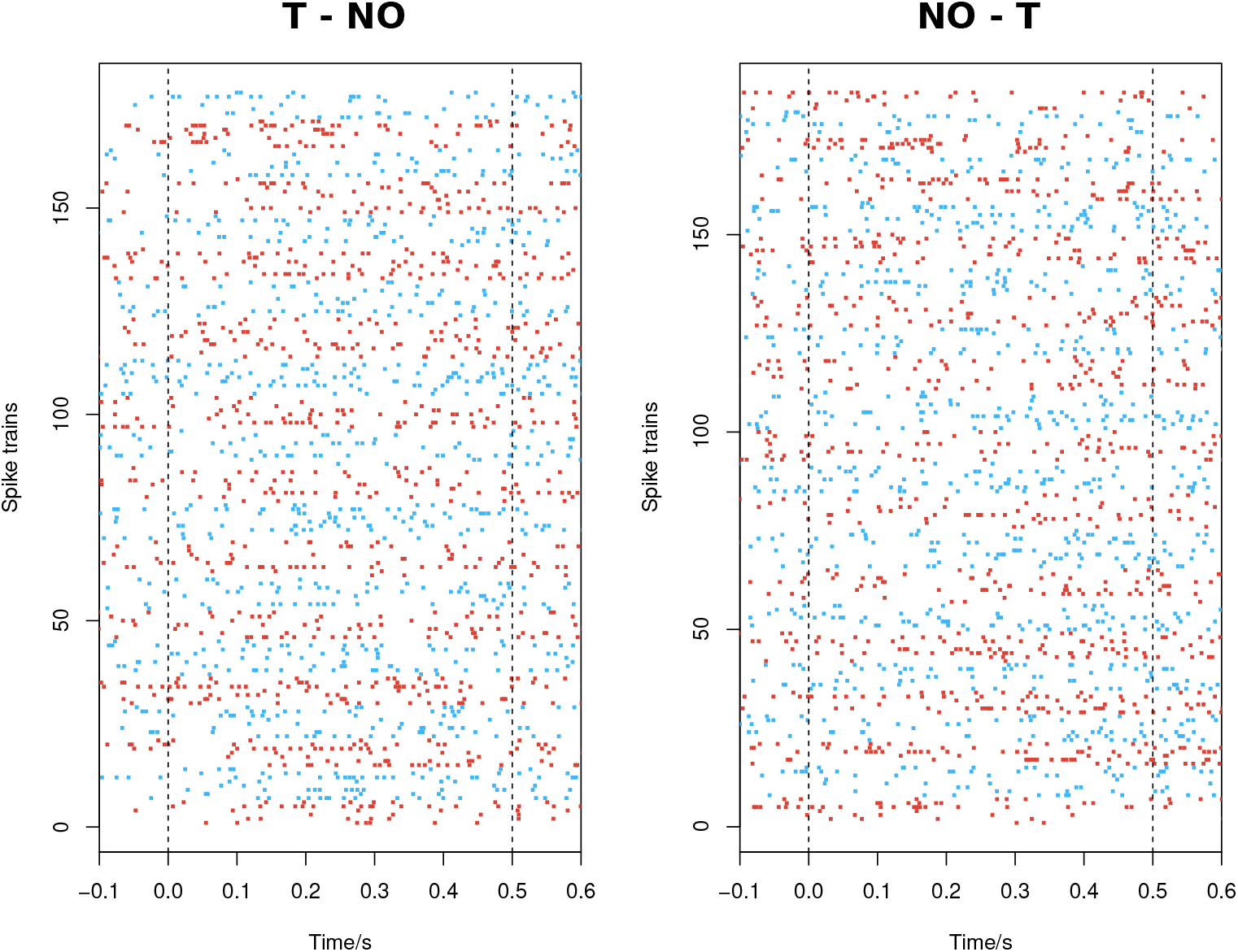
Spike trains of simultaneously recorded neurons. Session “MN110411” for two of the conditions. Each point in the figure denotes a spike at the time indicated by the *x*-axis. Different trials are presented alternately using the red and blue colors, and the simultaneously recorded spike trains within one trial are shown in the same color. The left and right panels show two different conditions. Dashed lines indicate the interval where the two stimuli are shown on the screen.

**S4 Fig.**
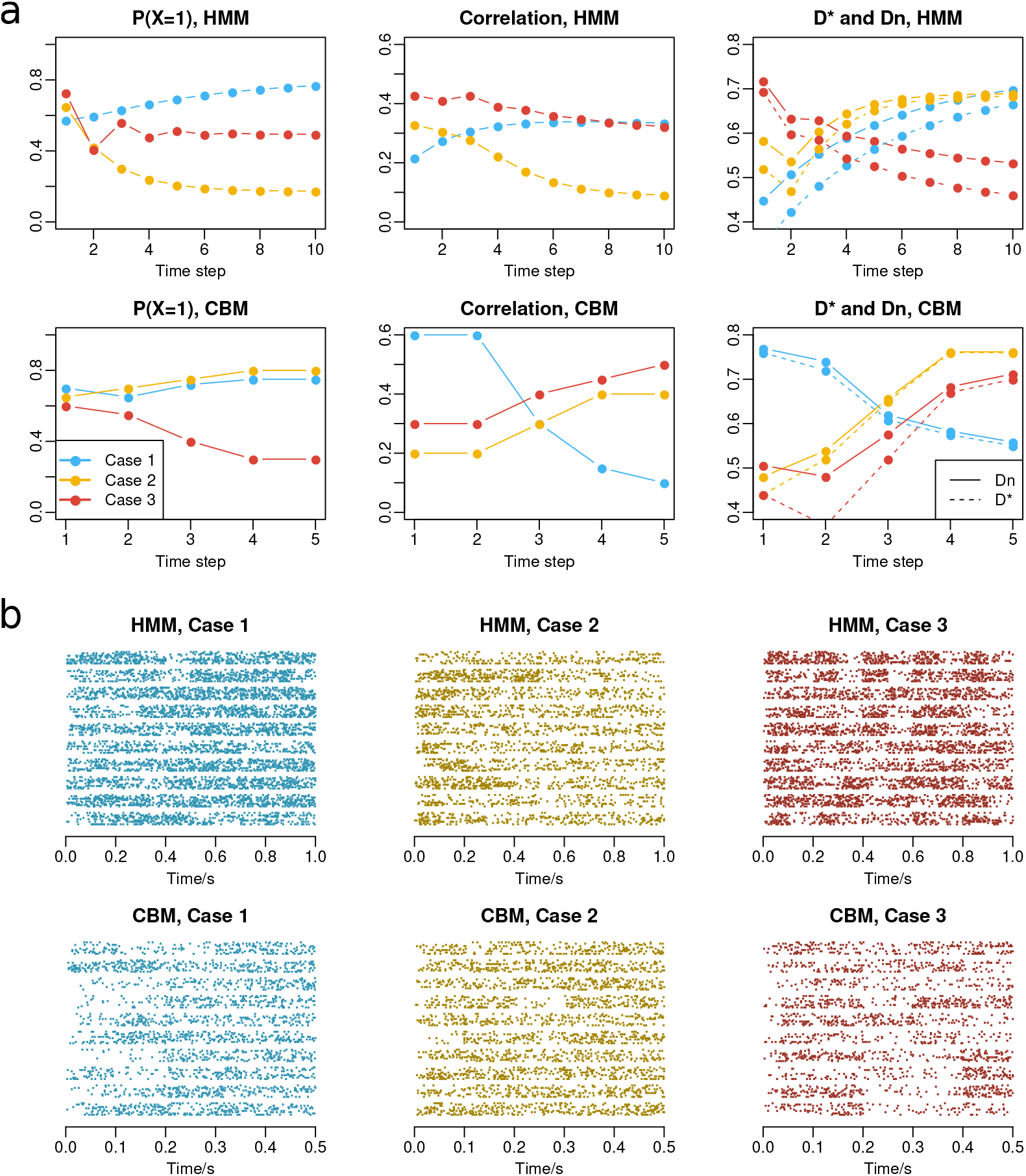
Examples of simulated data for the HMM and the CBM. a) The probabilities of attending the contralateral stimulus (left), the pairwise correlation coefficients (middle), and the deviation statistic values (right) are shown as functions of time. Different colors represent the three parameter settings. b) Example spike trains are shown for the corresponding case and model. In each sub-figure, 10 trials are shown, separated by horizontal white space lines. In each trial, 10 simultaneous spike trains are plotted.

**S5 Fig.**
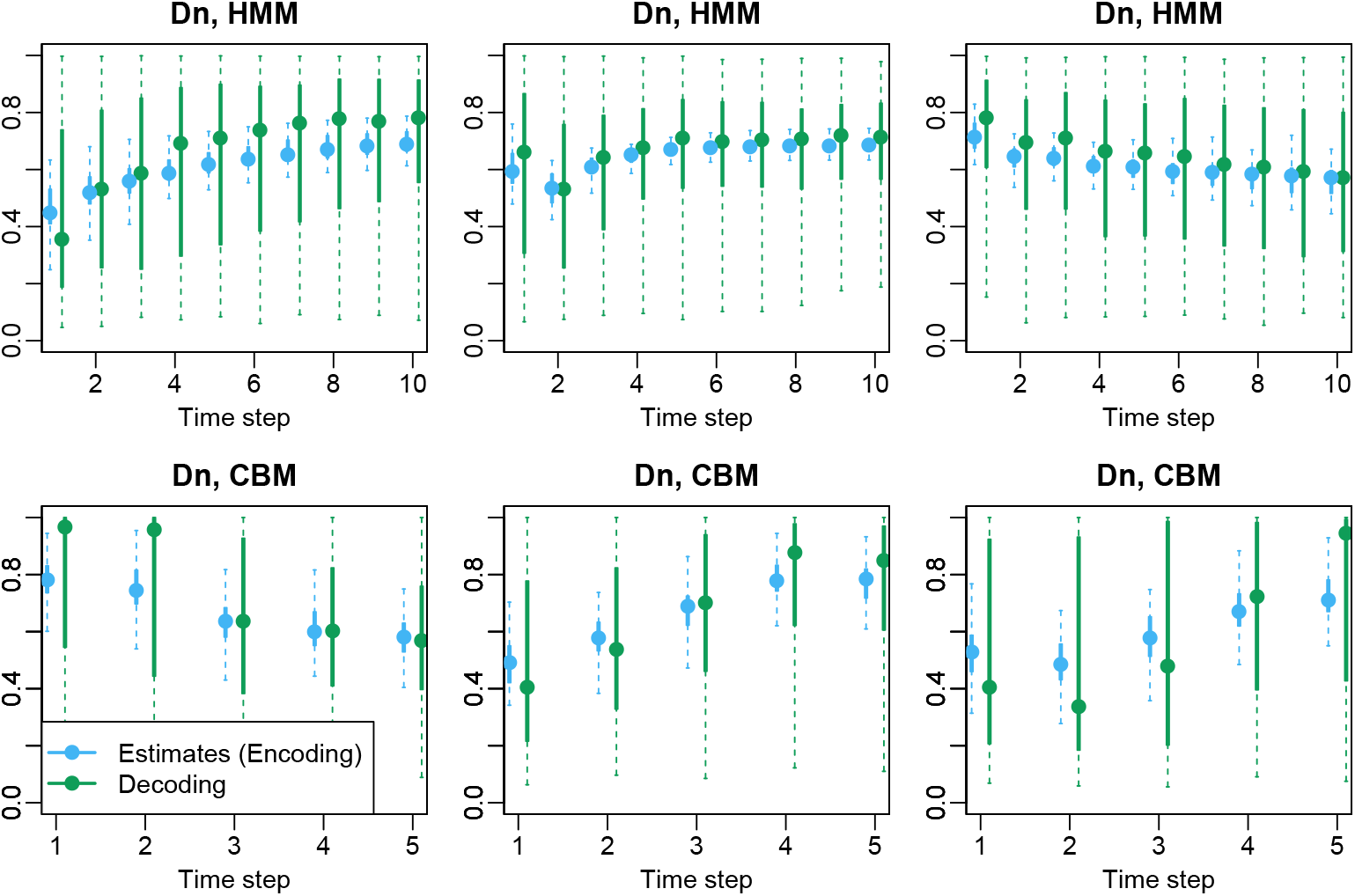
Deviation statistic values *D_n_* obtained from decoding or encoding analysis. Values are calculated from estimates (encoding, blue) or from decoding analysis (green) and are shown as quantiles of the 100 repetitions. The dashed lines represent the full 0 – 100% quantiles, and the solid lines represent the 25% – 75% quantiles. The dots are the medians.

**S6 Fig.**
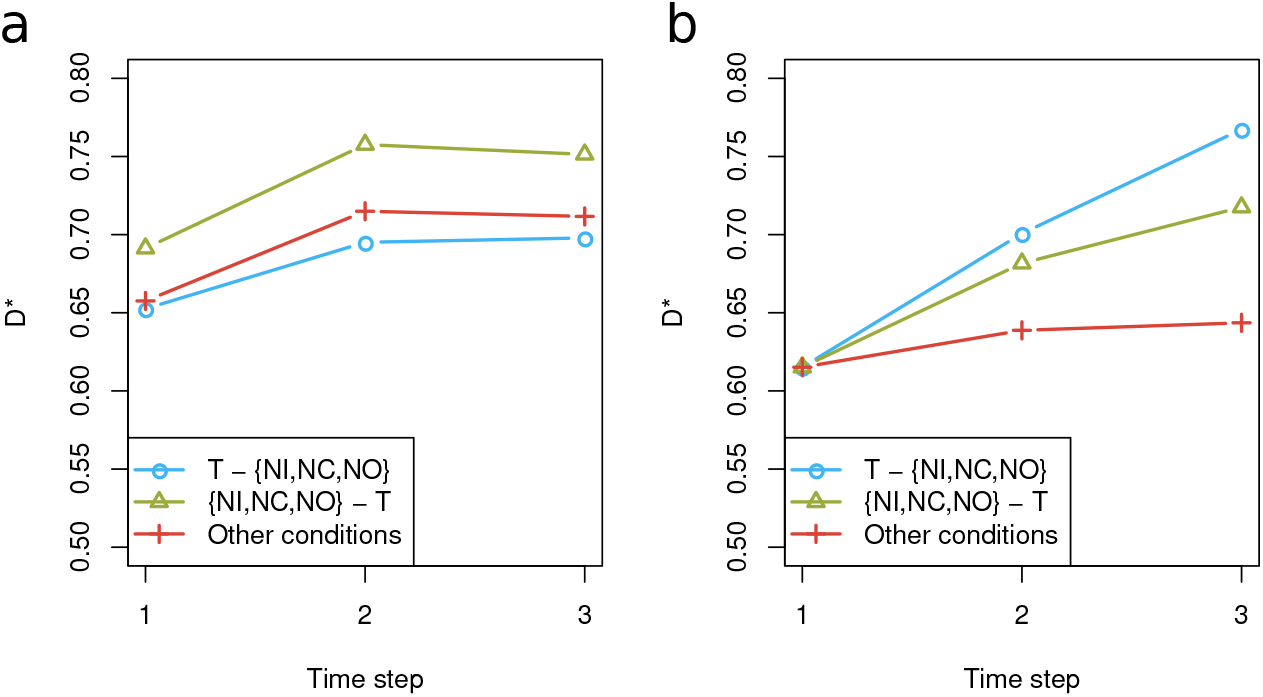
The *D*\* values allowing varying initial probabilities. These are from parameter estimates using two settings: assuming different initial probabilities for each condition (a), or fixing the same initial probabilities for all conditions as before (b).

